# Skin keratinocyte-derived SIRT1 and BDNF modulate mechanical allodynia in mouse models of diabetic neuropathy

**DOI:** 10.1101/2023.01.24.523981

**Authors:** Jennifer O’Brien, Peter Niehaus, Koping Chang, Juliana Remark, Joy Barrett, Abhishikta Dasgupta, Morayo Adenegan, Mohammad Salimian, Yanni Kevas, Krish Chandrasekaran, Tibor Kristian, Rajeshwari Chellappan, Samuel Rubin, Ashley Kiemen, Catherine Pei-Ju Lu, James W. Russell, Cheng-Ying Ho

**Author notes:** Correspondence to: Cheng-Ying Ho, Department of Pathology, Johns Hopkins University School of Medicine, 1800 Orleans St, M2101, Baltimore, MD 21287. These authors contributed equally to this work (Jennifer O’Brien, Peter Niehaus, Koping Chang).

## Abstract

Diabetic neuropathy is a debilitating disorder characterized by spontaneous and mechanical pain. The role of skin mechanoreceptors in the development of mechanical pain (allodynia) is unclear. We discovered that mice with diabetic neuropathy had decreased sirtuin 1 (SIRT1) deacetylase activity in foot skin, leading to reduced expression of brain-derived neurotrophic factor (BDNF) and subsequent loss of innervation in Meissner corpuscles, a mechanoreceptor expressing the BDNF receptor TrkB. When SIRT1 was depleted from skin, the mechanical allodynia worsened in diabetic neuropathy mice, likely due to retrograde degeneration of the Meissner-corpuscle innervating Aβ axons and aberrant formation of Meissner corpuscles which may have increased the mechanosensitivity. The same phenomenon was also noted in skin BDNF knockout mice. Furthermore, overexpression of SIRT1 in skin induced Meissner corpuscle reinnervation and regeneration, resulting in significant improvement of diabetic mechanical allodynia. Overall, the findings suggested that skin-derived SIRT1 and BDNF function in the same pathway in skin sensory apparatus regeneration and highlighted the potential of developing topical SIRT1-activating compounds as a novel treatment for diabetic mechanical allodynia.

## Introduction

Diabetic neuropathy is a common complication of diabetes and prediabetes, affecting up to 50% of the adult patients.^1^ Patients develop symptoms such as mechanical allodynia, spontaneous pain, numbness, paresthesia, and dysautonomia, with mechanical allodynia (neuropathic pain) reported in approximately 20% of the patients.^2^ Diabetic neuropathy, particularly the distal symmetric polyneuropathy subtype, is predominantly a small fiber neuropathy^3^ based on the clinical presentations and progressive loss of small nerve fibers seen on skin biopsies. Small fibers comprise of unmyelinated C fibers and small myelinated Aδ fibers^4^ and are responsible for sensory perceptions such as temperature and nociceptive pain.^5^ Large myelinated sensory fiber alterations have also been reported in early diabetic neuropathy ^6–8^, but are not well investigated. Among the sensory abnormalities experienced by diabetic neuropathy patients, burning and blunt pressure pain has been attributed to sensitization of C fibers, whereas pricking sensation is linked to Aδ fiber injuries.^9,10^ Mechanical allodynia, in comparison, is thought to be triggered by activation of Aß fibers.^11–13^ In the inflammatory pain mouse model, Aβ fibers appear to cause mechanical allodynia by switching their phenotype to one resembling nociceptive C fibers, thereby enhancing central sensitization to innocuous stimuli.^14^ A study by Xu et al. showed that blockade of Aß and not C fibers successfully inhibited mechanical allodynia, highlighting the potential of targeting Aß fibers as a novel therapeutic strategy for neuropathic pain.^15^

Aß mechanosensory neurons consist of slowly adapting (SA) and rapidly adapting (RA) low-threshold mechanoreceptors (LTMRs), which form sensory apparatus such as Merkel cell complexes, Ruffini endings, Meissner corpuscles and Pacinian corpuscles in glabrous (hairless) skin and longitudinal lanceolate endings in hairy skin.^16^ Among these Aß mechanosensory receptor end organs, Meissner corpuscles are the only one that possesses immunochemical properties similar to nociceptors^17^ in addition to their normal function of detecting gentle touch perception.^18^ Reduced densities of Meissner corpuscles have been observed in human patients with peripheral neuropathy such as chemotherapy-induced peripheral neuropathy^19^ and diabetic neuropathy.^20,21^ Creigh et al. used in-vivo reflectance confocal microscopy to visualize Meissner corpuscles in diabetic neuropathy patients, and detected decreased Meissner corpuscle densities early in the disease process. ^21^ The finding suggests that Meissner corpuscle alterations may be used as a marker to monitor disease progression or treatment response in diabetic neuropathy.

Meissner corpuscles are dually innervated by TrkB^+^ and Ret^+^ Aß mechanosensory afferents. A study by Neubarth et al. revealed loss of Meissner corpuscles and light touch perception in mice lacking TrkB in primary sensory neurons.^18^ Dhandapani et al. showed that photo-ablation of TrkB^+^ sensory neurons caused retraction of TrkB^+^ afferents from skin and alleviated mechanical allodynia in mouse models of neuropathic pain, suggesting that TrkB^+^ afferents are necessary and sufficient to convey mechanical allodynia after nerve injury.^22^ Neubarth et al. further demonstrated a dramatic reduction in the number of Meissner corpuscles upon elimination of brain-derived neurotrophic factor (BDNF), the TrkB ligand, from skin keratinocytes. The finding highlights the importance of skin-derived BDNF in early Meissner corpuscle development.^18^ To investigate whether skin-derived BDNF is also important for maintenance of mature Meissner corpuscles and for protection against diabetic neuropathy, we generated a conditional transgenic mouse that lacked BDNF expression in skin keratinocytes. We also created two skin-specific transgenic mouse lines that either lacked or overexpressed sirtuin 1 (SIRT1), a transcriptional activator of *BDNF*.

SIRT1 is an NAD^+^-dependent protein deacetylase which acts as a metabolism sensor in a variety of metabolic processes. Decreased activity or expression of SIRT1 has been observed in various tissues of diabetic patients possibly due to decreased systemic NAD^+^ biosynthesis^23–26^ or increased NAD^+^ degradation.^27,28^ SIRT1 deacetylates a number of proteins e.g. FoxO, p53 and PGC1α, and regulates transcription of their downstream targets.^29^ It has also been shown to upregulate brain BDNF expression to mediate neuroprotection against neurodegenerative disease.^30,31^ Through a diverse approach, we hereby investigated how alterations in skin keratinocyte-derived SIRT1 and BDNF cause injuries to Meissner corpuscles and their innervating Aβ nerve fibers in mouse models of diabetic mechanical allodynia.

There have been multiple animal models for diabetic mechanical allodynia. Popular models include streptozotocin (STZ)-induced rodent models for type 1 diabetes^32,33^ and high-fat diet (HFD) feeding mouse models for type 2 diabetes.^34,35^ The STZ models recapitulate the metabolic phenotypes of type 1 diabetes; however, they only develop mild mechanical allodynia.^33^ In comparison, the HFD models demonstrate well characterized mechanical pain phenotypes but are mainly pre-diabetic. Another rodent model that has been used to study mechanical allodynia in type 2 diabetes is the HFD combined with low-dose STZ model.^36,37^ This model is truly diabetic and may show more pronounced pain phenotypes compared with the HFD only model. For this study, we utilized both the HFD only as well as the HFD + STZ mouse model, both of which demonstrate robust peripheral neuropathy phenotypes and can be used to model early- and late-stage diabetic neuropathy, respectively.

## Materials and methods

### Animals and treatment

Animals were maintained and handled following the protocols approved by the University of Maryland Baltimore and Johns Hopkins University Animal Care and Use Committee. The C57BL/6J, *K5-CreER^T2^* (JAX 029155), *SIRT1^flo/flox^* (JAX 008041)^38^, *BDNF^flox/flox^*(JAX 033689)^39^, *K5-rtTA* (JAX 017579) and Ai14 *ROSA26 ^LSL-tdTomato^* (JAX 007914)^40^ mouse strains were acquired from the Jackson Laboratory. Generation of *TREbi-mSIRT1OE/mito-eYFP* was described previously.^41^ Any strain that did not develop impaired glucose tolerance was backcrossed with C57BL/6J for at least 6 generations. Cre-mediated recombination was induced in mice 3-7 months of age by intraperitoneal injection of 2 mg/day tamoxifen (Sigma-Aldrich) diluted with 300 μl corn oil (Sigma-Aldrich) for 3-5 consecutive days. Tamoxifen was given every 2-3 months to maintain Cre expression. Littermates were used to compare the phenotypic differences between the controls and KO/OE mice.

To create HFD only models, mice 3-7 months of age were fed with a HFD containing 36% fat (60% calories from fat), 20.5% protein (15% calories from protein), and 37.5% carbohydrate (26% calories from carbohydrate) (Bio-Serv, F3282). For KO mice, diet modification was initiated one week after tamoxifen injection. For OE mice, doxycycline (DOX) (Sigma-Aldrich) was added to the HFD at a dose of 200 mg/kg. To create HFD combined with low-dose STZ models, after 2-3 months of receiving HFD the mice were given 75 mg/kg STZ (Sigma-Aldrich) in 0.1M sodium citrate buffer, pH 5.0, through intraperitoneal injection, followed by a second dose of STZ at 50 mg/kg 3 days later.^36,37^ HFD continued for another month. Mice with random blood glucose ≥ 250 mg/dl^36,37^ or fasting blood glucose ≥ 200 mg/dl were considered diabetic.^42,43^

### Human subjects

Images of de-identified sural nerve biopsies previously collected for diagnostic purposes at University of Maryland Medical Center were the source of material for the study. The biopsies met the following selection criteria: non-neuropathic control or a clinical diagnosis of diabetic neuropathy. No protected health information was collected. The study belonged to federally defined Institutional Review Board-Exempt Research Category 4: Secondary Research or Existing Data, which includes “pathological specimens with information recorded by the investigator in such a manner that subjects cannot be identified, directly or through identifiers linked to the subjects”.

### Study design

To ensure that our results are robust, unbiased, and reproducible, the following was performed for all experiments: *1)* random assignment of mice; *2)* performance of behavioral studies by personnel blinded to the identity of the groups; *3)* blind assessment of outcome measures; *4)* specific exclusion criteria identified prior to initiation of all experiments; and *5)* assessment of sex. This study employed unique mouse models to address specific research questions. All experiments used littermate controls. To determine the number of mice needed, power analysis was performed to account for group and sex differences. The estimates of the sample sizes were based on performing Student’s t-test, 1-way or 2-way ANOVA, two tailed with α< 0.05 and a β=0.2 (80% power).

### Immunohistochemistry and quantification of intraepidermal nerve fiber density (IENFD)

Mouse hind paws were fixed in Zamboni’s fixative for 24-48 hours at 4°C and processed as previously described.^44^ 50 μm floating tissue sections were stained with the primary antibody rabbit anti-protein-gene-product 9.5 (PGP 9.5) (Dako, Z5116, 1:500) overnight and the secondary biotinylated goat anti-rabbit IgG antibody (Vector Laboratories, BA-1000-1.5) for 1 hour, followed by colorimetric detection using the VECTASTAIN ABC-HRP Kit (Vector Laboratories, PK-4000). Quantification of IENFD was performed blindly following the protocol described by Lauria et al.^44^

### Immunofluorescence staining

Tissue sections were obtained from mouse hind paw skin fixed and embedded following the IENFD protocol. For processing of spinal cord and DRG, mice were first perfused with 4% paraformaldehyde (PFA) in PBS. The nervous tissue was post-fixed in 4% PFA in PBS overnight and cryoprotected in 30% sucrose in PBS before being embedded in Tissue-Tek O.C.T. compound (Sakura Finetek). 30 μm tissue sections were incubated with blocking solution (5% normal goat serum in PBST [0.1% Triton X-100 in PBS]) at room temperature for 30 minutes and then with the primary antibody at 4 °C overnight. After being washed with PBST, tissue sections were incubated with the fluorophore-conjugated secondary antibody at room temperature for 1 hour, washed, stained with DAPI Solution (ThermoFisher, #62248, 1:5000), and mounted with Fluoromount-G (Southern Biotech). Images were acquired by a Carl Zeiss LSM700 or Leica 3i Spinning Disk confocal microscope. Primary antibodies used include rabbit anti-S100 beta (Proteintech, 15146-1-AP, 1:200), rabbit anti-S100 (Agilent, GA504), chicken anti-neurofilament heavy chain (NFH) (Abcam, ab4680, 1:50 or Aves Labs, 1:50), rabbit anti-DsRed (Takara, #632496, 1:500) and chicken anti-beta-tubulin 3 (Aves Labs, TUJ-0020, 1:500).

### Quantification of Meissner corpuscle and epidermal A**b** fiber density

Confocal S100/DAPI or NFH/DAPI images were acquired from 6-8 x 30 μm hind paw skin sections containing glabrous pads for each mouse and quantified blindly. Meissner corpuscle (epidermal Aβ fiber) density was defined by the total number of Meissner corpuscles (epidermal Aβ fiber) divided by the total area (total length of the foot pad bumps measured at the base of the epidermis x total thickness of the skin sections).^18^

### Three-dimensional (3D) reconstruction of mouse foot tissues

Meissner corpuscles were mapped in the foot skin of 9 mice: 3 controls, 3 mice fed with a HFD for 4 months, and 3 mice fed with a HFD for 3 months, treated with low-dose STZ and followed by continued HFD for 6 additional months. Samples were collected, paraformaldehyde-fixed, OCT embedded, and 75 serial histological sections were obtained for each sample. Serial sections were stained according to the scheme: (1) hematoxylin and eosin (H&E), (2) PGP9.5, (3) S100, repeat. Sections were digitized, and the serial images were reconstructed into digital tissue volumes using CODA, a novel technique for 3D mapping of serial histological images.^45^ Using CODA, six microanatomical structures were recognized in the H&E images: epidermis, vasculature, stratum corneum, extra cellular matrix, oil/sweat glands, and muscle, with a testing accuracy of 96.3%. Meissner corpuscles were mapped through quantification of positive antibody signal in the IHC images near the epidermal surface, with a testing accuracy of 94.8%. 3D renderings were created to visualize the foot microanatomy.

### 10x single cell RNA sequencing

Mouse paws were collected as biopsies from 4 wild-type (WT) control and 4 WT HFD mice or 2 *SIRT1* control (*SIRT1^flox/flox^*) and 2 *SIRT1* KO mice. The skin tissue was digested with 1000 U/ml collagenase and 300 U/ml hyaluronidase (Sigma-Aldrich) for 1.5 hour at 37 °C and 0.25% trypsin-EDTA (GIBCO) was added for additional digestion for 10 min. Digested tissues were suspended and washed with PBS containing 4% of fetal bovine serum, then filtered through 40 μm cell strainers to make single-cell suspensions. Samples were washed three times and centrifuged at 300 g at 4 °C for 10 min. Cell Hashing was used to label samples from distinct samples.^46^ Single cell suspensions were labeled with the TotalSeq Hashtag Antibodies (BioLegend) for 30 min on ice. Single cell suspensions were submitted to the Genome Technology Core facility at New York University Langone Medical Center and subjected to 10x genomics sequencing. Analysis of the data was performed using the Seurat V3 package.^47^

### Western blot analysis

Mouse foot skin was pulverized in liquid nitrogen and sonicated in RIPA lysis buffer (Sigma-Aldrich) containing protease inhibitor cocktail (Sigma-Aldrich, P8340). The lysates were centrifuged at 12,000 g at 4 °C for 10 min. 10 or 20 μg supernatants were analyzed by Western blot. The band intensity was normalized to GAPDH or vinculin. Primary antibodies used include rabbit anti-BDNF 19-HCLC (ThermoFisher, #710306, 1:500), rabbit anti-SIRT1 (Millipore, 07-131, 1:1000), rabbit anti-acetyl FOXO1 Lys294 (ThermoFisher, PA5-154560, 1:500), mouse anti-FOXO1 3B6 (ThermoFisher, MA5-17078, 1:500), and rabbit anti-vinculin (Cell Signaling Technology, #4650, 1:500).

### SIRT1 activity assay

Mouse foot skin was pulverized in liquid nitrogen and sonicated in lysis buffer (20 mM Tris-HCl, pH 7.5, 250 mM NaCl, 1 mM EDTA, 1% Triton X-100). The lysates were centrifuged at 12,000 g at 4 °C for 10 min. 5 ul supernatant was used for the SIRT1 activity assay following the manufacturer’s instruction (Abcam). Fluorescence intensity was normalized to the total protein concentration measured by Pierce BCA Protein Assay (ThermoFisher).

### Electron microscopy

Mouse foot skin tissue was fixed in 4% formaldehyde and 1% glutaraldehyde in phosphate buffer, and processed as previously described ^48^.

### Quantification of A**β** axons in human sural nerve biopsies

Toluidine blue stained semithin sections of sural nerve were imaged using a light microscope. Large myelinated (Aβ) axons (diameter > 5 μm) were blindly quantified, and their density was determined as the number of Aβ axons divided by the area of the nerve fascicle.

### Quantification of degenerated A**β** axons in subcutaneous nerve bundles in mouse foot skin

Mouse foot skin tissue was imaged by electron microscopy. Degenerated large myelinated (Aβ) axons (diameter > 5 μm) in subcutaneous nerve bundles were blindly quantified, and demonstrated as a percentage of the total Aβ axons.

### Behavioral testing

Static mechanical allodynia was assessed using von Frey monofilaments (0.02-4 g). Animals were placed on an elevated mesh grid. The test was initiated with a 2 g filament applied gently on the left mid hind paw until the filament started to bend and maintained for ∼2 s. A withdrawal response was considered valid only if the hind paw was completely removed from the platform. It was repeated 9 more times at 5-second intervals. The 50% withdrawal threshold was determined using the up-down method of Dixon.^41,49^

Dynamic mechanical allodynia was assessed using the dynamic brush assay.^50^ The adhesive removal (sticky tape) assay, light touch sensitivity assay, Hargreaves test, hot plate assay, acetone evaporation assay and pinprick test were performed following methods previously described by Duan et al.^50^ All behavior assays were assessed blindly.

### Sensory nerve conduction studies

Sensory NCV were measured from the tail using platinum electrodes, placed adjacent to the nerve, using a 60-80 mA square wave stimulus for 0.1-0.3 msec to obtain near nerve recordings. Orthodromic sensory tail NCV was obtained by placing the G1 (active) electrode at the base of the tail and stimulating 4 cm distally. Sensory responses were averaged from 10 trials until the nerve action potential response was stable. Tail temperature was maintained at 32-37 °C. The onset latency and peak amplitude were measured.^41^

### Statistical analysis

Statistical analyses were performed by unpaired two-tailed t test (one-tailed for Western blot due to small numbers of samples), one-way or two-way ANOVA with Tukey multiple comparison’s test. Simple linear regression analysis was performed by Prism software, version 10 (GraphPad). A *p* value of less than 0.05 was considered significant.

## Results

### Loss of subepidermal Aβ fibers and Meissner corpuscles a better indicator of diabetic mechanical allodynia

The two mouse models of diabetic neuropathy used in this study were high-fat diet (HFD) only and HFD combined with low-dose streptozotocin (STZ).^36,37^ For the HFD only model, mice were given a HFD for 3-4 months; for the HFD combined with STZ model, mice were given a HFD for 2-3 months, followed by 2 low doses of STZ and continued HFD for another month (Supplementary Fig. 1A, B). After 3-4 months of diet modification/STZ induction, both mouse models developed impaired glucose tolerance (Supplementary Fig. 1C-F). The HFD with low-dose STZ model developed both static and dynamic allodynia (Supplementary Fig. 1I, J), whereas the majority of the HFD only mice demonstrated static but not dynamic allodynia (Supplementary Fig. 1G, H). While some HFD only mice also developed dynamic allodynia, the result was not statistically significant compared to their baseline (Supplementary Fig. 1H).

Upon examination of the sensory apparatus involved in pain sensation in foot skin, both mouse models of diabetic neuropathy demonstrated a profound loss of Meissner corpuscles compared to the control mice (Fig. 1A, C). The Meissner corpuscle loss was accompanied by degeneration or retraction of their innervating Aβ cutaneous sensory axons (Fig. 1B, D). While the intraepidermal nerve fiber densities (IENFD), which measure predominantly the free nerve endings, were also lower in the diabetic neuropathy mice than the control mice, the difference was not as obvious (Fig. 1A, E). Of note, when we measured the IENFD on PGP9.5 stained sections, we counted only the small nerve fibers and meticulously avoided counting the large nerve fibers innervating the Meissner corpuscles. Overall, diabetic neuropathy caused 50-80% loss of Meissner corpuscles and Aβ as opposed to 30% loss of free nerve endings in foot skin of the mice. Simple linear regression analysis (Fig. 1F) further supported that subepidermal Aβ fiber density is a better predictor of mechanical allodynia than IENFD for both mouse models of diabetic neuropathy. Based on the phenotypic characterization, we believe that the two mouse models of diabetic neuropathy are similar, with the HFD + STZ model demonstrating more severe and advanced neuropathy phenotypes.

**Fig. 1.**
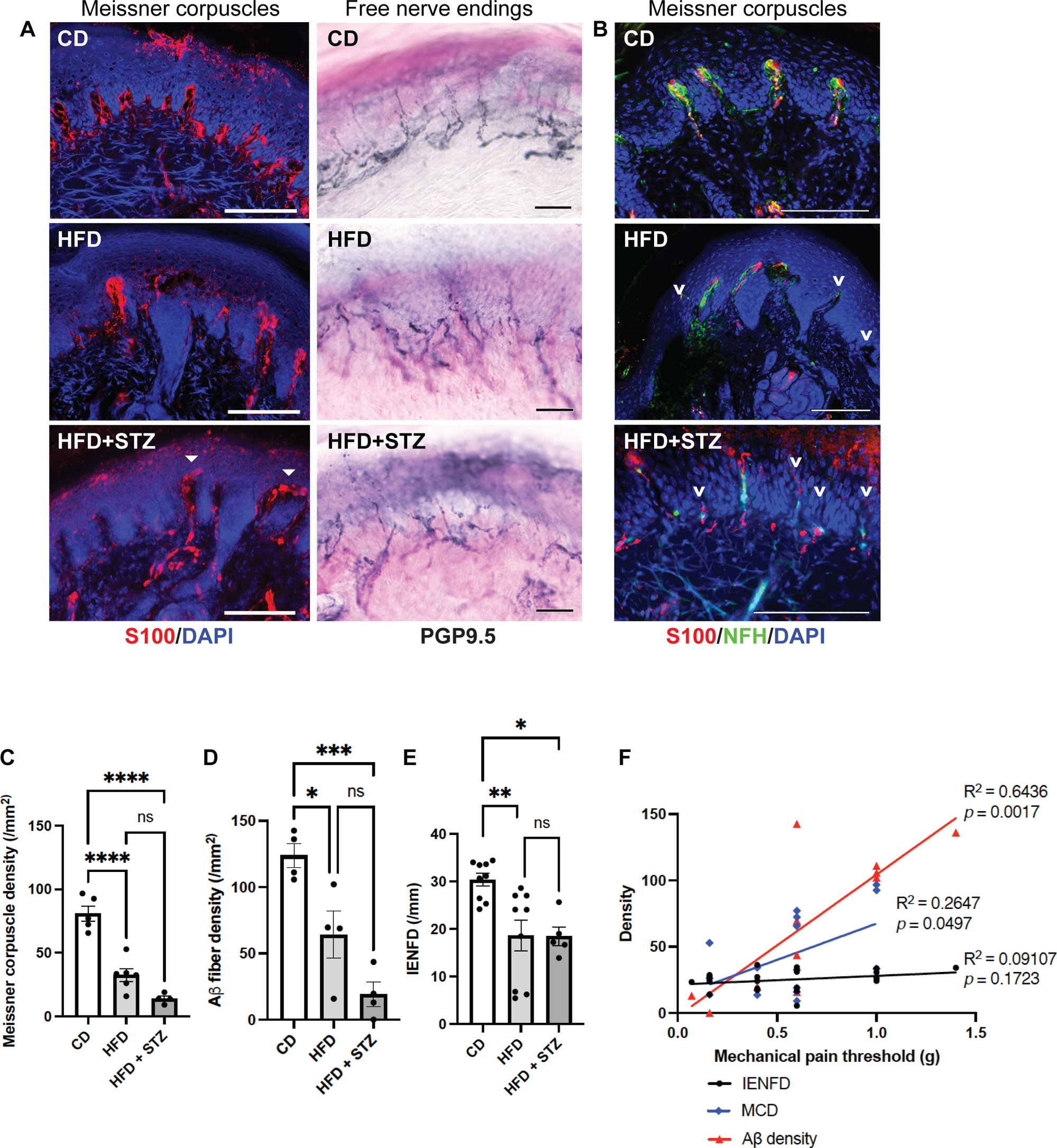
Subepidermal Aβ fiber density a better indicator of diabetic mechanical allodynia. (**A)** Immunofluorescence (IF) images of Meissner corpuscles and immunohistochemistry (IHC) images of free nerve endings in foot skin from control diet (CD) mice and two mouse models of diabetic neuropathy: high-fat diet (HFD) mice and HFD mice treated with low-dose streptozotocin (HFD + STZ). Meissner corpuscles are the S100^+^ rod-shaped terminations present in the dermal papillae near the dermal-epidermal junction. Filled arrowheads indicate the aberrant horizontally elongated corpuscles occasionally noted in HFD + STZ mice. (**B)** IF images of Meissner corpuscles double stained with S100 and neurofilament heavy-chain (NFH). Arrowheads indicate Aβ afferents of Meissner corpuscles undergoing axon degeneration or retraction. **(C)** Meissner corpuscle, **(D)** Subepidermal Aβ fiber and **(E)** free nerve ending densities (intraepidermal nerve fiber densities [IENFD]) in foot skin from CD, HFD and HFD + STZ mice. Statistics were performed by one-way analysis of variance (ANOVA). **(F)** Simple linear regression analysis. Meissner corpuscle density (MCD): n = 13; Aβ fiber density: n = 12; IENFD: n = 22. Scale bars represent 100 μm in IF and 50 μm in IHC images. Error bars indicate standard error of means (SEM). Statistics were performed by one-way analysis of variance (ANOVA): **** *p* < 0.0001, *** *p* < 0.001, ** *p* < 0.01, * *p* < 0.05, ns = non-significant.

To further investigate the spatial and structural changes of Meissner corpuscles, free nerve endings and their innervation in the dermis, we utilized a cutting-edge deep learning-based imaging platform named CODA.^45,51^ Developed by Kiemen and colleagues, CODA is a method that enables three-dimensional (3D) reconstruction of formalin-fixed, serially sectioned histological samples for study of large volumes of tissue. CODA uses image registration to create digital volumes from serial tissue images and deep learning semantic segmentation^52^ to label components in hematoxylin & eosin (H&E) and immunohistochemistry images. This novel technique allows us to study the anatomic relationship between sensory innervation and the adjacent skin structures, and to evaluate whether alterations of sensory innervation occur focally or globally.

The 3D model of foot skin demonstrated reduced numbers of Meissner corpuscles and free nerve endings in mice with diabetic neuropathy (Supplementary Fig. 2). In the advanced neuropathy model for which mice were continued on HFD for an extended period of time (a total of 9 months) after the STZ treatment, Meissner corpuscles (S100^+^/ PGP9.5^+)^ were completely absent while few free nerve endings (PGP9.5^+^) remained. In comparison, mice that were on HFD only for 4 months had a more modest loss in Meissner corpuscles and free nerve endings. Notably, not only sensory apparatus in epidermis was lost, their innervation in deep dermis also vanished, suggesting that retrograde degeneration of sensory axons occurs in the advanced stage of diabetic neuropathy. The finding may explain why patients with late-stage diabetic neuropathy often experience sensory loss.

### Decreased expression of BDNF in foot skin of mice with diabetic neuropathy

To identify molecular events that led to loss of Meissner corpuscles and their innervation in mice with diabetic neuropathy, we performed single-cell RNA-sequencing (scRNA-seq) on foot pad skin of mice fed with a HFD for 4 months and age matched controls fed with a standard chow diet. The analyses focused on keratinocytes which were the most abundant cells in foot pad skin. Six major cell types were identified after unbiased clustering analysis (Fig. 2A). Overall, there were no global changes in cell type proportions or gene expression landscape in the foot skin of HFD mice (Fig. 2B). We then focused on the expression of the skin-derived neurotrophic factors, which are known to regulate the development or survival of specific cutaneous sensory afferents.^53^ In mouse foot skin, the most abundant neurotrophic factor was neurturin (NRTN), followed by BDNF. The remaining skin-derived neurotrophic factors were expressed in < 1% of the foot skin cells (Supplementary Fig. S3A), and therefore may be less important in supporting cutaneous afferents. NRTN is a member of the glial cell line-derived neurotrophic factor (GDNF)-family ligands. Mice lacking NRTN demonstrate loss of P2X3^+^ epidermal afferents due to depletion of GFRα2^+^ neurons in dorsal root ganglia (DRG)^53,54^, whereas mice lacking cutaneous BDNF expression demonstrate loss of Aδ longitudinal lanceolate endings in hairy skin and Meissner corpuscles in glabrous skin.^18,55^ In the mouse foot pad where Meissner corpuscles reside, BDNF was expressed in all layers of epidermis with higher expression in basal and spinous keratinocytes (Fig. 2C). In foot skin of HFD mice, the number of keratinocytes expressing BDNF as well as the expression level were both decreased in epidermal keratinocytes compared to the control mice (Fig. 2D). The finding was further confirmed by the Western blot analysis, as the protein level of BDNF was also decreased in the foot skin of HFD mice (Fig. 2E, F). We speculated that downregulation of BDNF expression in skin of HFD mice was due to decreased enzymatic activity or decreased expression of SIRT1^56–59^, a protein deacetylase regulating BDNF transcription through the TORC1 and CREB pathway.^30^ Further assessment of SIRT1 deacetylase activity by fluorometric assay and by Western blotting analyzing the acetylation status of its substrate FoxO1 both revealed decreased deacetylase activity in the foot skin of HFD mice (Fig. 2G-I). SIRT1 expression levels in foot skin, however, were similar in HFD and control mice (Fig. 2H, J). Interestingly, the pattern of SIRT1-expressing cells in foot skin (Fig. 2K and Supplementary Fig. 3A) was nearly identical to that of BDNF-expressing cells (Fig. 2C and Supplementary Fig. 3A). The similarity in epidermal localization further suggests that the two skin-derived molecules may have a functional relationship in supporting mature Meissner corpuscles and protecting them against diabetes-induced damages.

**Fig. 2.**
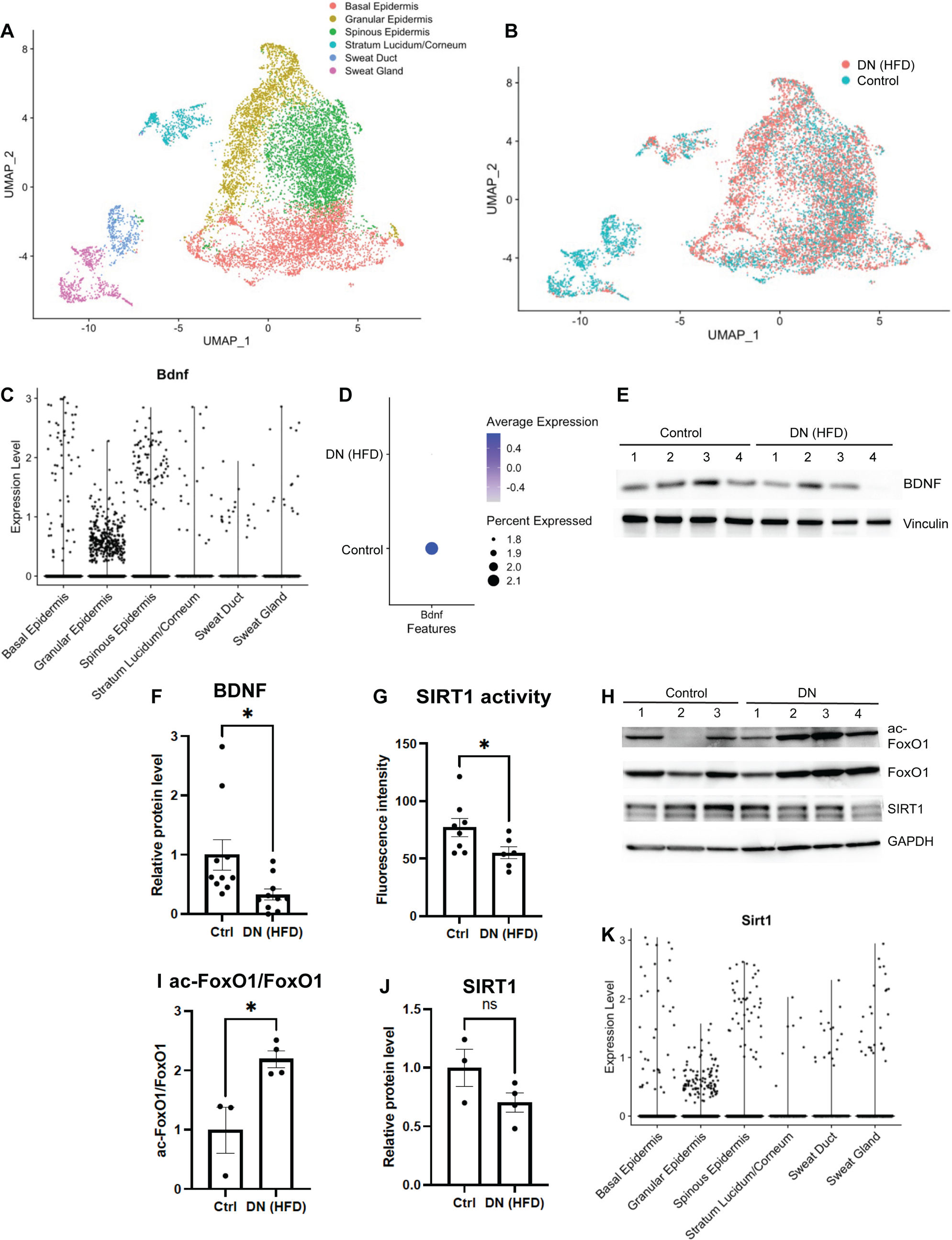
Decreased expression of BDNF in foot skin of mice with diabetic neuropathy. **(A)** UMAP projection of single-cell RNA-seq (scRNA-seq) data from mouse foot pad skin epithelial cells colored by cell subtype annotation (n = 11,079). **(B)** UMAP plot of foot pad skin epithelial cells from mice with diabetic neuropathy (DN) (HFD for 4 months) and control mice (n = 4 for each group). **(C)** Violin plot demonstrating BDNF expression levels in different subtypes of epithelial cells in mouse foot pad skin. **(D)** Dot plot indicating the differential BDNF expression in epidermal keratinocytes between the control and DN mice. The size of the circles represents the percentage of BDNF-expressing cells, and the color of the circles represents the level of BDNF expression. Note that there is no BDNF expression (circle) in the intra-pad epidermis. **(E)** A representative Western blot of mouse foot skin lysates with antibodies against BDNF and vinculin, a loading control. **(F)** Comparison of normalized mature BDNF protein level in foot skin of control and DN mice (n = 9 for each group). **(G)** Measurement of SIRT1 deacetylase activity in mouse foot skin by fluorometric assay (n = 8 for the control; n = 6 for DN). **(H)** Western blot probing for the acetylated form of the SIRT1 substrate FoxO1. Also shown are blots for total FoxO1, SIRT1 and GAPDH (loading control), respectively. **(I)** Comparison of acetyl-FoxO1/FoxO1 protein ratio and **(J)** normalized SIRT1 protein level in foot skin of control (n = 3) and DN mice (n = 4). **(J)** Violin plot demonstrating SIRT1 expression levels in different subtypes of epithelial cells in mouse foot pad skin. Error bars indicate SEM. Statistics were performed by Student’s t-test: * *p* < 0.05, ns = non-significant.

### Diabetic mechanical allodynia exacerbated by depletion of the BDNF transcriptional regulator SIRT1 from skin

To elucidate the role of skin-derived SIRT1 in the pathogenesis of diabetic neruopathy, we generated a conditional keratinocyte-specific *SIRT1* knockout (KO) strain *Keratin 5 (K5)-CreER^T2^*;*SIRT1^flox/flox^*. Tamoxifen was given to adult mice at least 3 months of age to bypass the potential effect of SIRT1 depletion on development of sensory axons. Tamoxifen treatment induced Cre-mediated recombination in basal keratinocytes and produced a non-functional truncated SIRT1 protein due to in-frame deletion of exon 4 (Supplementary Fig. 4A).^38^ We noticed leaky Cre-mediated expression in basal keratinocytes at approximately half efficiency of the expression induced by full-dose tamoxifen (Supplementary Fig. 4B). *SIRT1^flox/flox^*was therefore included as an additional control (Control 1) besides *K5-CreER^T2^*;*SIRT1^flox/flox^* without tamoxifen (Control 2). K5-Cre-mediated gene recombination, however, was tissue-specific, as expression of the reporter gene was not detected in DRG or spinal cord (Supplementary Fig. 4C).

Skin keratinocyte-specific *SIRT1* KO mice appeared to be grossly phenotypically normal, except for patchy back hair loss in < 10% of the mice. They did not develop significant mechanical allodynia until after receiving 3 months of HFD: dynamic allodynia in *SIRT1* KO mice was more severe than both control groups, whereas static allodynia was only significantly worse than Control 1 (Fig. 3A, B). Notably, while approximately 2/3 of the keratinocyte-specific *SIRT1* KO mice demonstrated static allodynia, the remaining 1/3 were hyposensitive to pain after diet modification (Fig. 3C). Given that hypoalgesia or sensory loss is a feature of late-stage diabetic neuropathy^33^, keratinocyte-specific *SIRT1* KO may be used to model advanced-stage disease. Progression of allodynia was monitored for 7 months after the initiation of HFD. In keratinocyte-specific *SIRT1* KO mice, both static and dynamic allodynia worsened over time and peaked at 5 months of diet modification (Fig. 3D, E). Note that Control 2 mice also developed static and dynamic allodynia but to a lesser degree possibly due to leaky expression of Cre recombinase (Supplementary Fig. 4B). The finding suggests that depletion of even just 30-50% of the SIRT1 from skin keratinocytes may be sufficient to worsen diabetic mechanical allodynia.

**Fig. 3.**
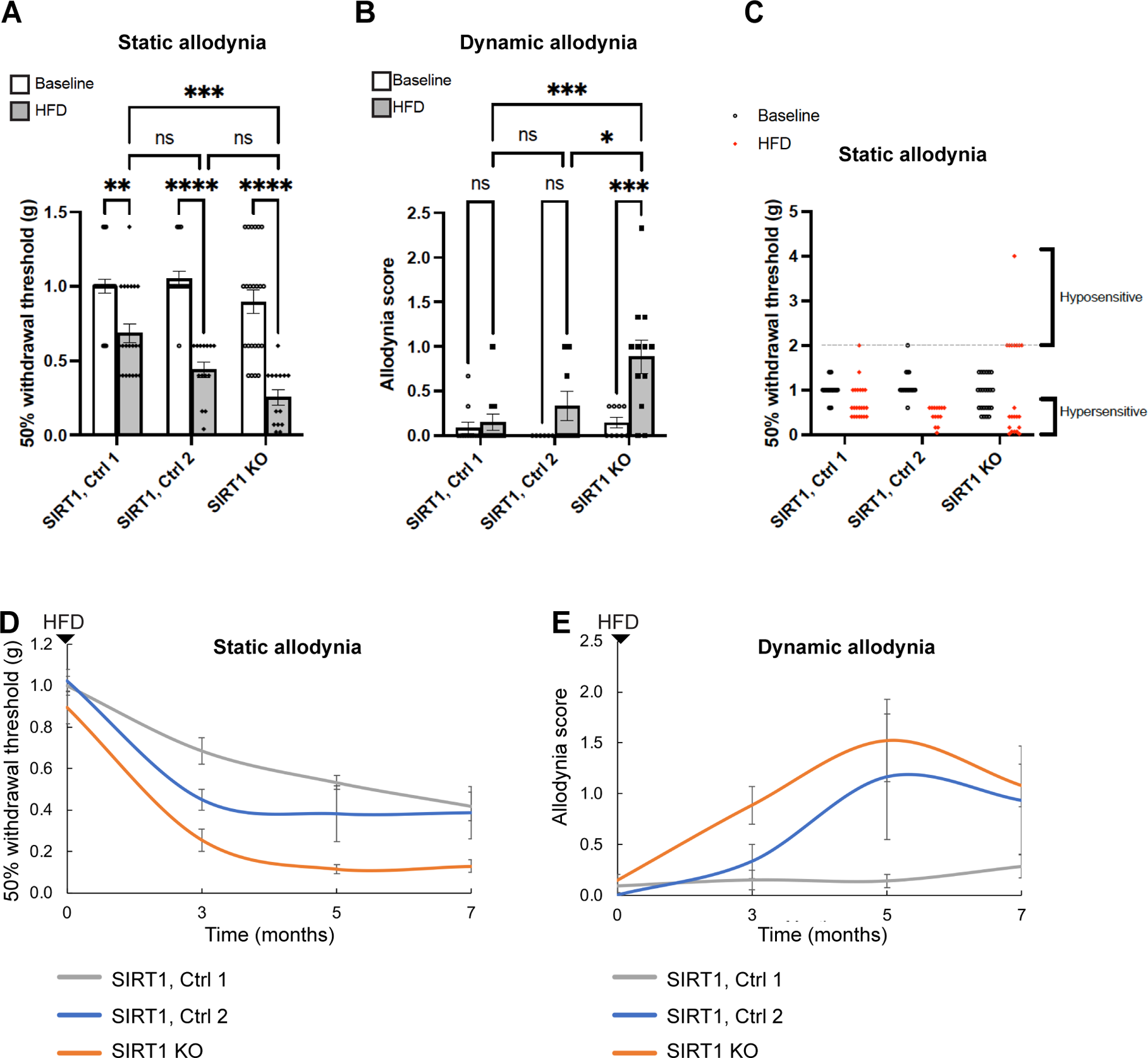
Exacerbated diabetic mechanical allodynia in adult mice lacking skin keratinocyte-derived SIRT1. **(A)** Assessment of static allodynia in keratinocyte-specific *SIRT1* KO mice and two control groups: Control 1: *SIRT1^flox/flox^* and Control 2: *K5-CreER^T2^*;*SIRT1^flox/flox^*receiving no tamoxifen but showing leaky Cre expression (**Supplementary Fig. 4**). Static allodynia was assessed by the von Frey assay. Note that only mice with normal (1 or 1.4 g) or decreased pain thresholds (< 1 g) were charted. **(B)** Assessment of dynamic allodynia in *SIRT1* KO and control mice. Dynamic allodynia was assessed by the dynamic brush assay. **(C)** Mice with increased pain thresholds (≥ 2g) are charted with the rest of the mice. **(D)** Progression of static and **(E)** dynamic allodynia in *SIRT1* KO and control mice. Error bars indicate SEM. Statistics were performed by two-way ANOVA: **** *p* < 0.0001, *** *p* < 0.001, ** *p* < 0.01, * *p* < 0.05, ns = non-significant.

The mechanical allodynia appeared to be the only sensory domain in foot (glabrous) skin affected by depletion of skin keratinocyte-derived SIRT1. There was no noticeable tactile hyposensitivity, thermal hyperalgesia, cold allodynia or abnormal nociception in *SIRT1* KO mice (Fig. 4). Depletion of keratinocyte-derived SIRT1 also did not exacerbate HFD-induced glucose metabolism defects (Supplementary Fig. 5). The findings further supported that skin-derived SIRT1 has a major and specific role in the pathogenesis of diabetic mechanical allodynia.

**Fig. 4.**
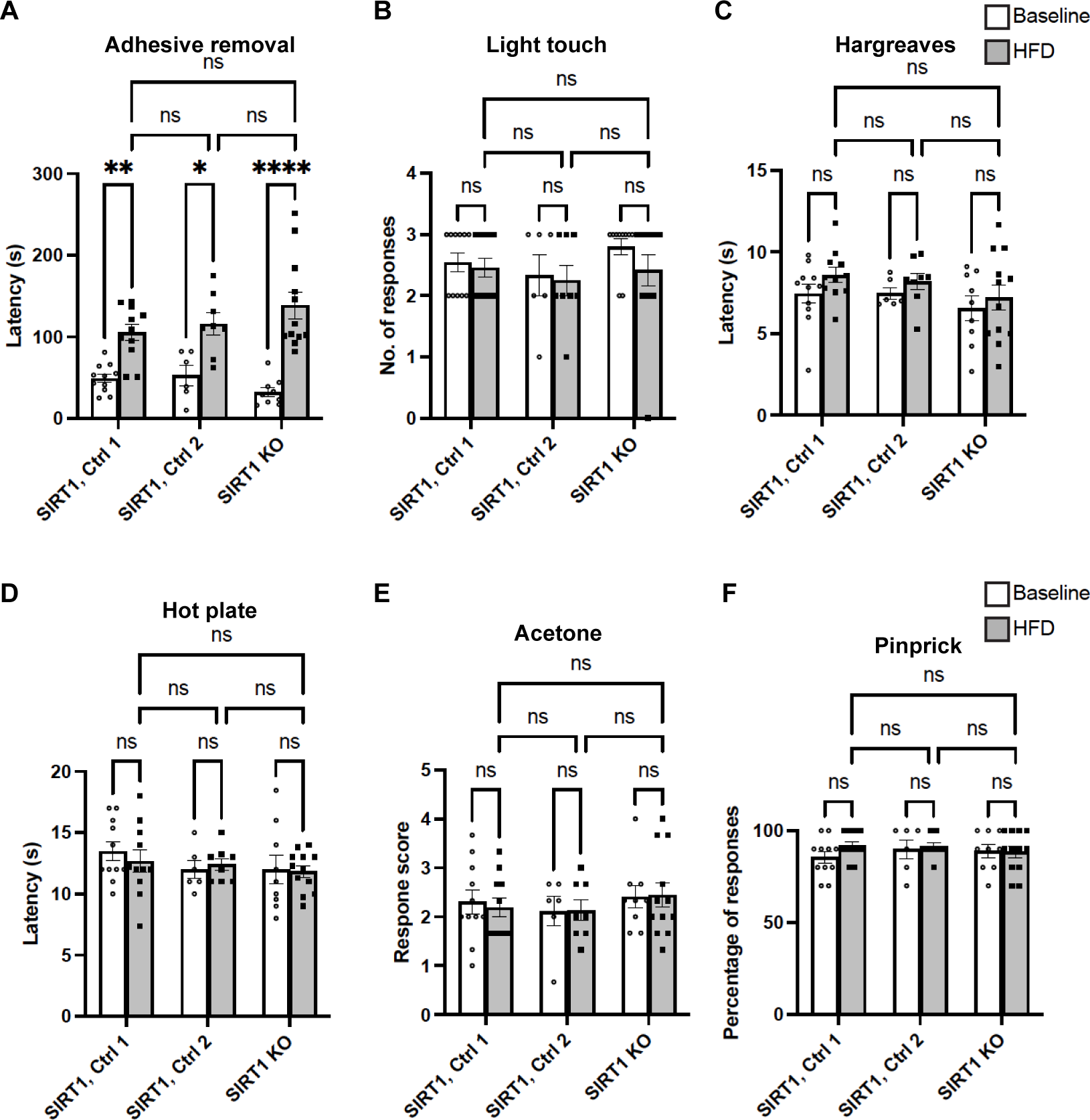
Largely normal non-allodynia sensory domains of foot (glabrous) skin from mice lacking skin keratinocyte-derived SIRT1. **(A)** Adhesive removal assay, **(B)** Light touch assay, **(C)** Hargreaves assay, **(D)** Hot plate assay, **(E)** Acetone evaporation assay, and **(F)** Pinprick assay. Note that in all 3 groups the latency to detection of the adhesive was increased after HFD treatment, but there was no significant difference between the controls and *SIRT1* KO mice. Error bars indicate SEM. Statistics were performed by two-way ANOVA: **** *p* < 0.0001, ** *p* < 0.01, * *p* < 0.05, ns = non-significant.

### Worsened Meissner corpuscle loss along with large sensory axon injury in skin *SIRT1* KO mice

After 3 months of HFD, Meissner corpuscle densities were markedly reduced in keratinocyte-specific *SIRT1* KO when compared to Control 1. The change was also observed in Control 2 mice but to a lesser degree and not statistically significant (Fig. 5A, B). Loss of Meissner corpuscles was accompanied by loss of their innervating Aβ axons (Fig. 5B, D). Free nerve endings, in comparison, were not affected by depletion of SIRT1 from keratinocytes, as the KO and control mice had similar IENFD (Fig. 5A, C). In addition to Meissner corpuscle loss, mice lacking keratinocyte-derived SIRT1 also demonstrated aberrant Meissner corpuscle morphology. A fraction of the Meissner corpuscles had the tip of the corpuscle misshaped into a long horizontal bar (Fig. 5F, G). Notably, the horizontal bar contained only the glial cell lamellae but no large axon terminals (Fig. 5H, I). Although it remains unclear how these aberrant Meissner corpuscles are related to an increase in mechanical pain, one possible mechanism is that the horizontally extended tip increases the area of contact with the stimulus and therefore enhances the skin mechanosensitivity.

**Fig. 5.**
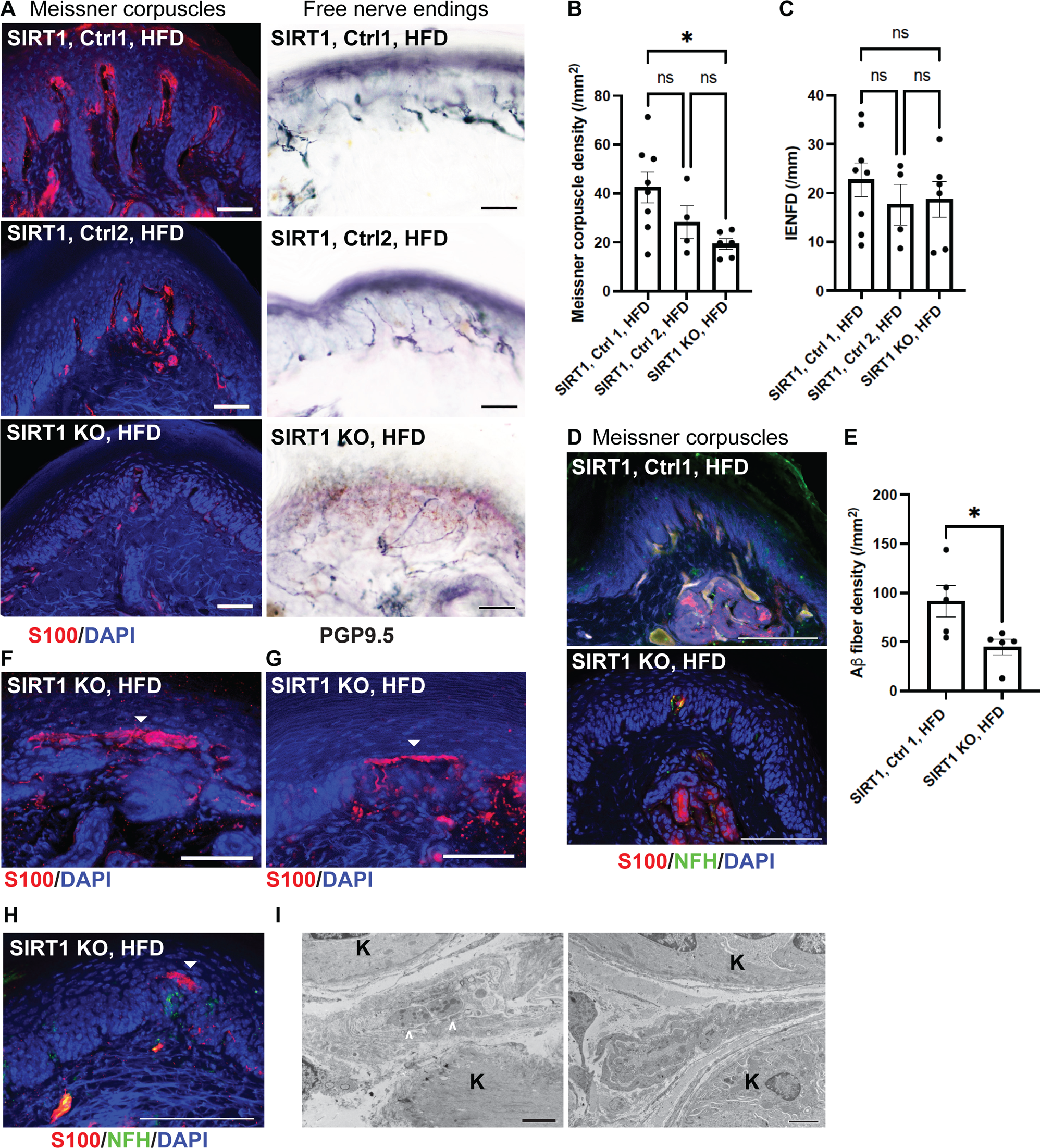
Reduced numbers and abnormal morphology of Meissner corpuscles in foot skin from mice lacking skin keratinocyte-derived SIRT1. **(A)** IF images of Meissner corpuscles and IHC images of free nerve endings in foot skin from keratinocyte-specific *SIRT1* KO, Control 1 and Control 2 mice. Scale bars represent 50 μm. **(B)** IF images of Meissner corpuscles double stained with S100 and NFH. **(C)** Meissner corpuscle, **(D)** free nerve ending (IENFD) and **(E)** epidermal Aβ fiber densities in foot skin from *SIRT1* KO and control mice. **(F-H)** IF images showing aberrant Meissner corpuscles with the tip appearing as a long horizontal bar in epidermis (filled arrowheads). Note in **(H)** that the aberrant tip did not contain the innervating NFH^+^ large axons. Scale bars represent 50 μm. Scale bars represent 100 μm for b, e-g. **(I)** Electron micrographs demonstrating a relatively normal Meissner corpuscle (left) containing nerve endings (arrowheads) ensheathed by glial cell lamellae and an aberrant Meissner corpuscle (right) with elongated glial cell lamellae but no nerve endings. K = keratinocytes. Scale bar represents 2 μm. Error bars indicate SEM. Statistics were performed by one-way ANOVA: * *p* < 0.05, ns = non-significant.

When Meissner corpuscle-innervating large myelinated (Aß) sensory axons were traced further back, severe axon degeneration was noted in subcutaneous nerve bundles of foot skin from keratinocyte-specific *SIRT1* KO mice (Fig. 6A). The percentage of degenerated Aß sensory axons was significantly higher in *SIRT1* KO mice than the control. Since Meissner corpuscles are innervated by Aß axons, the finding suggests that depletion of skin keratinocyte-derived SIRT1 may cause retrograde degeneration of Aß mechanosensory axons. The notion was further supported by decreased sensory nerve conduction velocities (NCV) (Fig. 6C), which mostly reflects abnormalities in large sensory fibers. Changes in large myelinated sensory axons were also observed in human patients with advanced diabetic neuropathy: sural nerve biopsies showed an average of 57% reduction in large myelinated axons in diabetic neuropathy patients compared to control individuals (Fig. 6D, E). Taken together, the keratinocyte-specific *SIRT1* KO mice nicely recapitulate the features of advanced diabetic neuropathy which are lacking in most animal models, and may be used to model late-stage painful diabetic neuropathy.

**Fig. 6.**
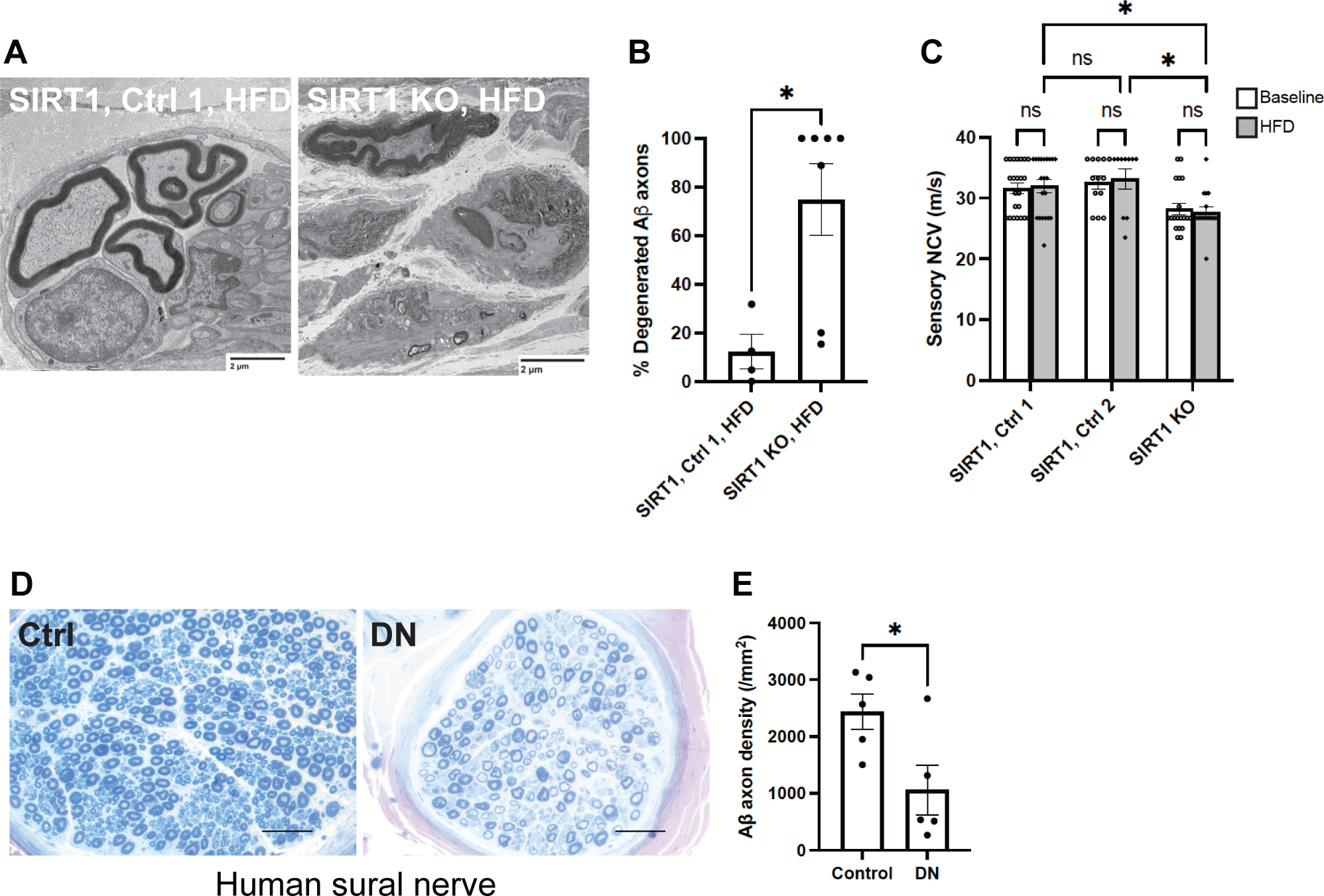
Large sensory (Aβ) fiber abnormalities and injuries in mice lacking skin keratinocyte-derived SIRT1 as well as human patients with diabetic neuropathy. (**A)** Electron micrographs of subcutaneous nerve bundles in foot skin from keratinocyte-specific *SIRT1* KO and control mice. **(B)** Quantification of degenerated large myelinated (Aβ) axons in subcutaneous nerve bundles (n = 4 for Control 1; n = 7 for SIRT1 KO). **(C)** Tail sensory nerve conduction studies of *SIRT1* KO, Control 1 and Control 2 mice. Statistics were performed by two-way ANOVA. **(D)** Sural nerve biopsies from human diabetic neuropathy (DN) and control patients. (**E)** Quantification of Aß axons in sural nerve. Error bars indicate SEM. Statistics were performed by Student’s t-test: * *p* < 0.05, ns = non-significant.

### Similar peripheral neuropathy phenotypes in skin BDNF and SIRT1 KO mice

scRNA-seq was performed on foot skin of keratinocyte-specific *SIRT1* KO and control mice to identify gene perturbations that might account for the exacerbated DN phenotypes associated with depletion of keratinocyte-derived SIRT1 (Supplementary Fig. 6A). The clustering analysis did not reveal any major changes in cell subpopulations or gene expression landscape (Supplementary Fig. 6B). However, targeted analysis of skin-derived neurotrophic factors showed down-regulation of BDNF in keratinocyte-specific *SIRT1* KO, whereas all the other major neurotrophic factors were in fact up-regulated (Fig. 7A). To further characterize the functional relationship between skin-derived BDNF and SIRT1, we generated a conditional keratinocyte-specific *BDNF* KO: *Keratin 5 (K5)-CreER^T2^*;*BDNF^flox/flox^*. Similar to keratinocyte-specific *SIRT1* KO, *BDNF* KO mice developed both static and dynamic allodynia after diet modification (Fig. 7B, C). In addition, their remarkable loss of Meissner corpuscle and epidermal Aβ fibers (Fig. 7D-F, H), occasional formation of aberrant Meissner corpuscles (Fig. 7I), and relatively unaffected free nerve endings (Fig. 7D, G) and glucose tolerance (Fig. 76A, B) all resembled the phenotypes of keratinocyte-specific *SIRT1* KO mice. The results indicate that BDNF and SIRT1 may operate in the same pathway to maintain the function and survival of Meissner corpuscles.

**Fig. 7.**
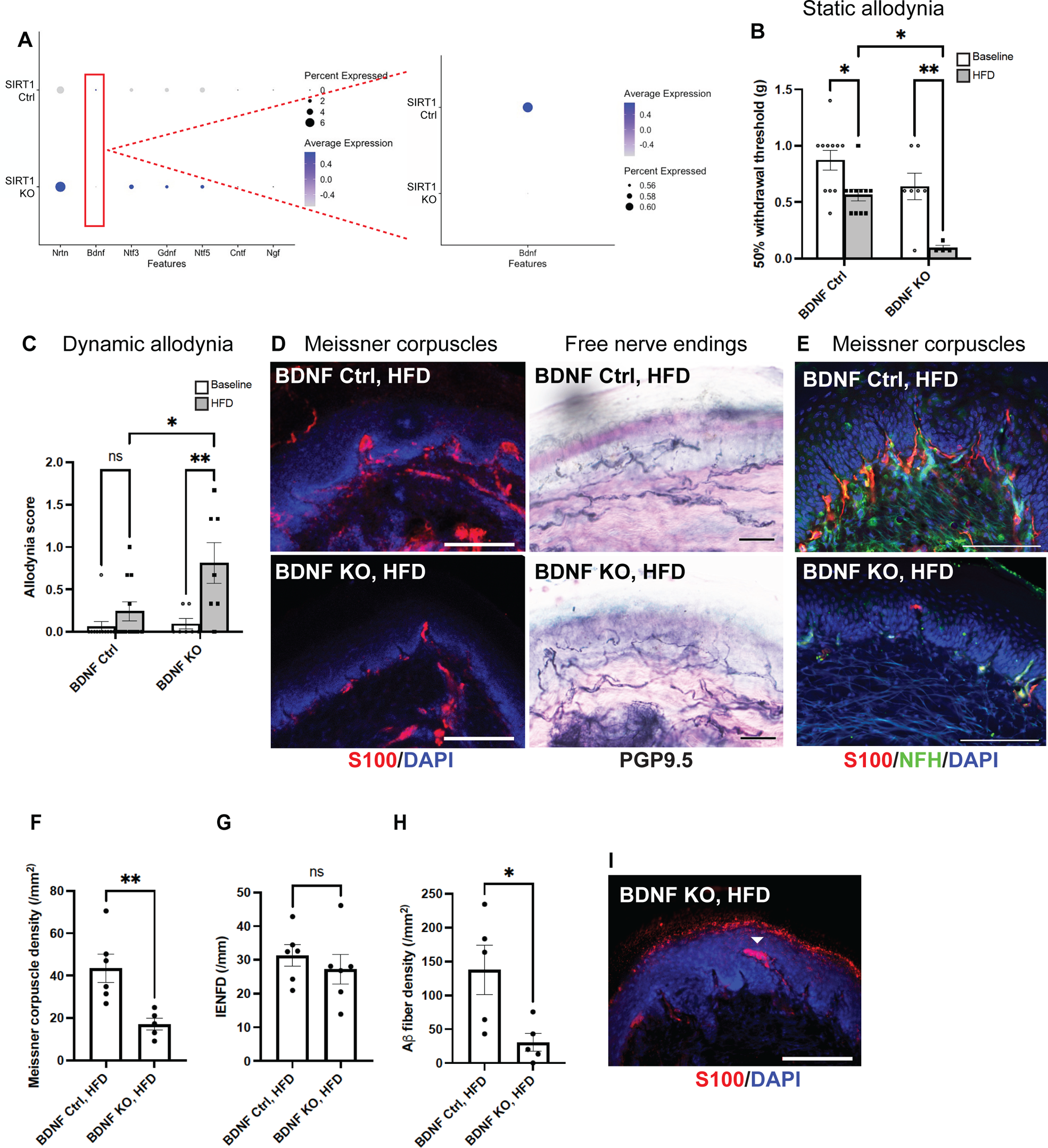
BDNF as the only major neurotrophic factor down-regulated in foot skin of mice lacking skin keratinocyte-derived SIRT1. **(A)** Dot plot based on scRNA-seq data **(Supplementary** Fig. 6) indicating the differential expression of neurotrophic factors in keratinocytes of foot skin between Control 1 (*SIRT1^flox/flox^*) and keratinocyte-specific *SIRT1* KO mice. The size of the circles represents the percentage of cells expressing a given neurotrophic factor, and the color of the circles represents the expression level of that neurotrophic factor. Similar to *SIRT1* KO, keratinocyte-specific *BDNF* KO developed more severe **(B)** static and **(C)** dynamic allodynia than the controls after 3 months of HFD. Statistics were performed by two-way ANOVA. **(D)** IF images of Meissner corpuscles and IHC images of free nerve endings in foot skin from *BDNF* KO and control mice. **(E)** IF images of Meissner corpuscles double stained with S100 and NFH. **(F)** Meissner corpuscle, **(G)** free nerve ending (IENFD) and **(H)** epidermal Aβ fiber densities in foot skin from *BDNF* KO and control mice. **(I)** Occasional Meissner corpuscle with abnormal morphology (filled arrowhead). Scale bars represent 100 μm for IF and 50 μm for IHC images. Error bars indicate SEM. Statistics were performed by Student’s t-test: ** *p* < 0.01, * *p* < 0.05, ns = non-significant.

### Diabetic neuropathy phenotypes reversed by overexpression of SIRT1 in skin

Since KO of keratinocyte-derived *SIRT1* worsened the diabetic neuropathy phenotypes, we generated an inducible transgenic mouse overexpressing keratinocyte-derived SIRT1 to evaluate whether diabetic neuropathy phenotypes could be alleviated. A Tet-On mouse strain keratinocyte-specific SIRT1OE was created by crossing *K5-rtTA*, a mouse harboring a Keratin 5 promoter-driven reverse tetracycline transactivator^60^, with *TREbi-mSIRT1OE/mito-eYFP*, a mouse expressing both mouse SIRT1 protein (mSIRT1) and mito-eYFP under the regulation of a tetracycline response element (TRE).^41^ Transgene expression can be induced and visualized when mice are fed with a doxycycline (DOX) diet (Fig. 8A). The mice were started on a HFD first, and then divided into two groups: the SIRT1OE group receiving HFD containing DOX and the control group receiving HFD only. Two months after the DOX treatment, there was significant improvement of diabetic mechanical allodynia, as the static allodynia thresholds of SIRT1OE mice returned to normal (Fig. 8B). Improvement of dynamic allodynia was also observed in SIRT1OE mice but not statistically significant due to the larger variation in allodynia scores in both groups (Fig. 8C). In addition, a complete recovery of Meissner corpuscles and Aβ fibers was noted (Fig. 8D-F, H). The mechanism of Meissner corpuscle and Aβ fiber recovery was most likely due to induction of BDNF expression in foot skin by SIRT1 overexpression (Fig. 8I-L). Notably, free nerve endings in foot skin were also increased by SIRT1 overexpression (Fig. 8D, G), while the glucose tolerance impairment was unchanged (Fig. S7C, D). The mechanism of increased innervation by free nerve endings, however, was unclear.

**Fig. 8.**
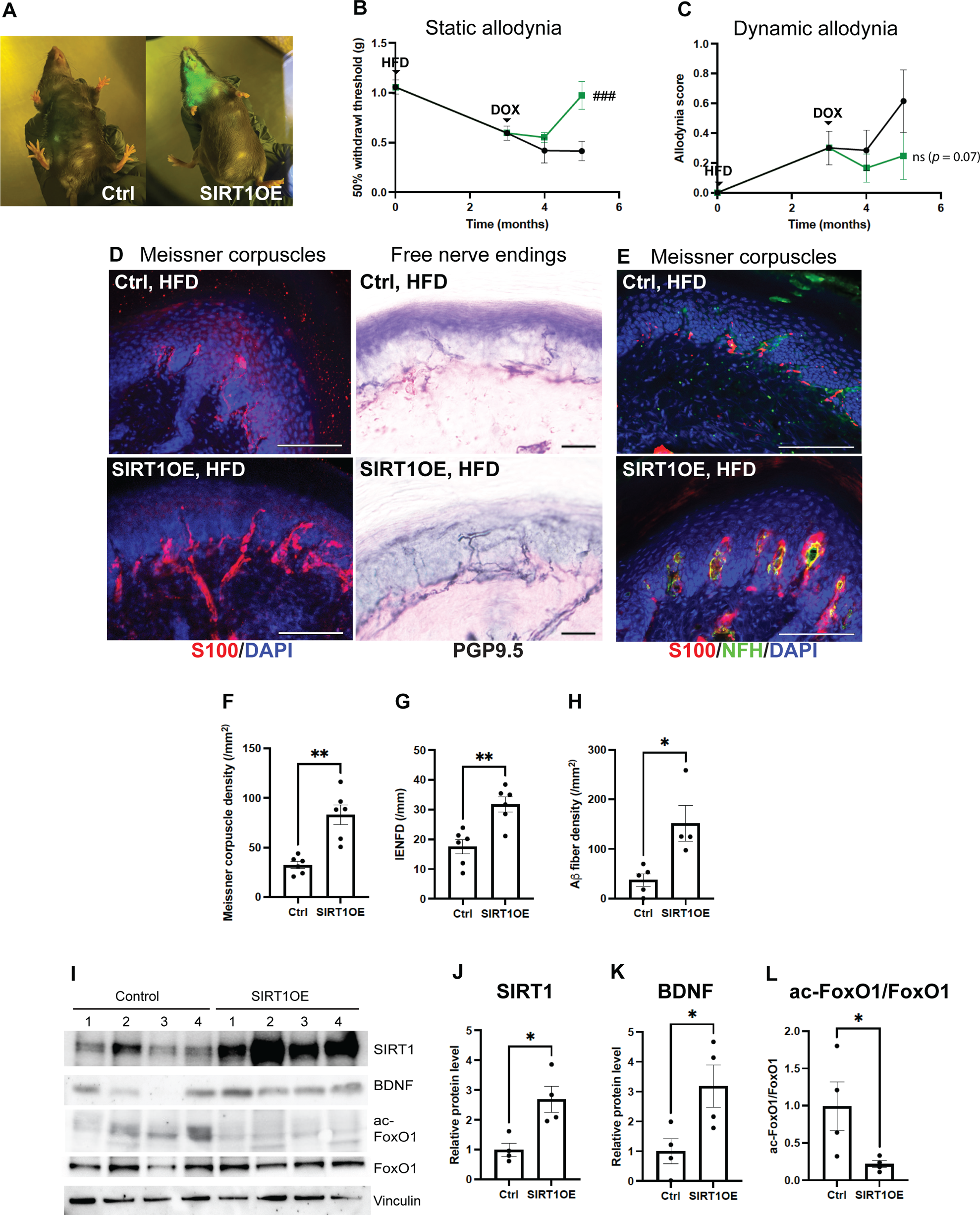
Diabetic neuropathy phenotypes rescued by overexpression of keratinocyte-derived SIRT1 (OE). **(A)** Skin of SIRT1OE mouse glowing in blue light due to simultaneous expression of a mito-eYFP transgene after one month of doxycycline (DOX) diet. **(B)** Diabetic static allodynia relieved after 2 months of DOX-induced SIRT1 overexpression in skin keratinocytes. Two-way ANOVA: ###: Control (n = 7) vs. SIRT1OE after 2 months of DOX (n = 7), *p* = 0.0003. **(C)** Improved diabetic dynamic allodynia but results not statistically significant. Two-way ANOVA: ns: Control (n = 7) vs. SIRT1OE after 2 months of DOX (n = 4), *p* = 0.07. **(D)** IF images of Meissner corpuscles and IHC images of free nerve endings in foot skin from skin SIRT1OE and control mice. **(E)** IF images of Meissner corpuscles double stained with S100 and NFH. **(F)** Meissner corpuscle, **(G)** free nerve ending (IENFD) and **(H)** epidermal Aβ fiber density in foot skin from SIRT1OE and control mice. **(I)** Western blot analysis of foot skin lysates from SIRT1OE (n = 4) and control mice (n = 4). Comparison of **(J)** normalized SIRT1 **(K)** normalized BDNF and **(L)** acetylated FoxO1 protein levels in foot skin of SIRT1OE and control mice. Scale bars represent 100 μm for IF and 50 μm for IHC images. Error bars indicate SEM. Statistics were performed by Student’s t-test: ** *p* < 0.01, * *p* < 0.05.

## Discussion

Diabetic neuropathy is a sensory disorder with pathology starting as loss of the skin sensory apparatus and gradually progressing to degeneration of all fiber types in the peripheral nerves. Patients present with painful sensation such as mechanical allodynia and thermal hyperalgesia first and eventually become insensate. Elucidating the molecular mechanisms underlying the pain and sensory disorders associated with diabetes requires preclinical studies in animal models. Existing animal models, however, have serious limitations. Most of the animal models only demonstrate mild neuropathy. Advanced phenotypes such as axonal degeneration and losses in the peripheral nerves are largely absent in these models.^61^ In addition, pain behavioral testing in rodents, such as the von Frey assay, can only measure the withdrawal thresholds and may not accurately recapitulate the clinical features of diabetic mechanical allodynia in human patients. By depleting SIRT1 and BDNF in the skin, our study created a novel mouse model of advanced diabetic neuropathy. This new model may provide insights into mechanisms of disease progression and facilitate the development of novel therapeutics for advanced neuropathy.

In addition to pain and sensory abnormalities, patients with diabetic neuropathy often have skin manifestations, such as diabetic foot ulcer and diabetic dermopathy.^62,63^ Cutaneous manifestations may be secondary to neuropathy or a predisposing factor to neuropathy. In the latter case, skin changes caused by diabetes may induce or accelerate the damage to the sensory apparatus, leading to peripheral neuropathy. Our study supports this notion by demonstrating that neuropathy is exacerbated by depletion of the skin-derived molecule SIRT1 or BDNF.

SIRT1 is a metabolic sensor, the activity of which is reduced in diabetes due to diminished levels of its activator NAD^+^.^23–28^ In addition to NAD^+^, SIRT1 activity is regulated by another metabolic sensor AMP-activated protein kinase (AMPK). AMPK is activated by changes in the AMP/ATP ratio and its dysregulation has been linked to diabetes and other metabolic disorders.^64^ Deficient AMPK signaling in diabetes may compromise SIRT1 signaling and lead to decreased SIRT1 target gene expression.^65^ This mechanism may also account for the downregulation of BDNF expression in diabetic skin along with the imbalanced NAD^+^ homeostasis.

Skin-derived BDNF plays a critical role in the early development of skin sensory apparatus, such as Aδ longitudinal lanceolate endings and Meissner corpuscles.^18,55^ Whether or not skin-derived BDNF remains essential for the maintenance of these sensory apparatus in adulthood is unknown. Data from our study suggest that skin-derived BDNF and its transcriptional regulator SIRT1 may not be required for the survival of the mature Meissner corpuscles under normal conditions, but their deficiency may precipitate the Meissner corpuscle loss and Aβ fiber degeneration associated with aging^66^ or diseases. In the sensory nervous system, BDNF is primarily secreted by DRG neurons. After peripheral nerve injury, BDNF has been shown to be upregulated in large-diameter DRG neurons and anterogradely transported to spinal dorsal horns.^67^ It is possible that skin keratinocyte-derived BDNF also anterogradely transports to large-diameter TrkB^+^ mechanosensory neurons to support their sensory axons. When there is decreased expression of skin-derived BDNF in the setting of diabetes as we demonstrated in this study, large sensory axons may degenerate due to the lack of trophic support.

While our data highlight the importance of Meissner corpuscles in mediating diabetic mechanical allodynia, it is unclear why loss of Meissner corpuscles or other skin sensory receptors in fact generates pain. This key question has been the subject of intensive research for many years but remains largely unanswered. Data from our study showed that not only Meissner corpuscles were diminished, but their Aß afferents also degenerated. Afferent injury may alter the expression or function of ion channels, leading to hyperexcitability of primary sensory neurons as a mechanism of neuropathic pain.^68^ In addition, our study demonstrated that mice with more severe diabetic mechanical allodynia, such as the keratinocyte-specific *SIRT1* and *BDNF* KO, appear to have aberrant Meissner corpuscle morphology: the top lamellar disks became elongated and extended horizontally instead of vertically to the epidermis (Fig. 5H, I and 7I). A study from Gangadharan et al. revealed that neuropathic pain can arise from peripheral miswiring, such as Meissner corpuscles reinnervated by nociceptors instead of LTMRs, after nerve injury.^69^ While we did not notice such findings in our models of diabetic neuropathy, the aberrant regeneration and reinnervation of Meissner corpuscles by Aβ fibers in our models may magnify the sensation by increasing the contact area with the touch stimulus, and may therefore represent a potentially novel mechanism of neuropathic pain.

Lastly, reorganization of neural circuits^70,71^ or neuroinflammation^72^ may occur in the spinal cord after peripheral nerve injury. Our current study has focused on peripheral sensitization but cannot exclude a role of central sensitization in the pathophysiology of diabetic mechanical allodynia. Further studies investigating the alterations in spinal circuits in the setting of diabetes are required to fully elucidate the mechanisms underlying diabetic mechanical allodynia.

In summary, our findings demonstrate the importance of the skin microenvironment, particularly the levels of skin keratinocyte-derived BDNF and its transcriptional regulator SIRT1, in the maintenance of Meissner corpuscle afferents and regulation of mechanical allodynia in animal models of diabetic neuropathy. Our results also highlight these skin-derived molecules as promising therapeutic targets for diabetic neuropathy given their potential for development of topical therapeutics.

## Data availability

scRNA-seq data will be deposited in NCBI Gene Expression Omnibus.

## Abbreviations

3D: 3-dimensional
BDNF: brain-derived neurotrophic factor
DOX: doxycycline
DRG: dorsal root ganglia
ER: estrogen receptor
GDNF: glial-derived neurotrophic factor
H&E: hematoxylin & eosin
HFD: high-fat diet
IENFD: intraepidermal nerve fiber density
K5: keratin 5
KO: knockout
LTMR: low-threshold mechanoreceptor
NAD: nicotinamide adenine dinucleotide
NCV: nerve conduction velocity
NFH: neurofilament heavy chain
NGF: nerve growth factor
NRTN: neurturin
NT: neurotrophin
OE: overexpression
PGP9.5: protein gene product 9.5
RA: rapidly-adapting
SA: slowly-adapting
scRNA-seq: single-cell RNA-sequencing
SIRT1: sirtuin 1
STZ: streptozotocin
TRE: tetracycline response element
YFP: yellow fluorescent protein

