## Supplemental Figures and Information for "Skin keratinocyte-derived SIRT1 and BDNF modulate mechanical allodynia in mouse models of diabetic neuropathy"

### Supplemental Figure Legends

**Supplementary Fig. 1. Two mouse models of diabetic neuropathy (DN) used in this study:** **(A)** High-fat diet (HFD) only mouse model; **(B)** HFD combined with low-dose streptozotocin (STZ) mouse model. **(C)** 2-hour intraperitoneal glucose tolerance tests (GTT) on mice receiving HFD only. (**D)** 2-hour GTT on mice receiving HFD with low-dose STZ. **(E)** The glucose area under the curve (AUC) from mice receiving HFD only. **(F)** The glucose AUC from mice receiving HFD with low-dose STZ. **(G)** Decreased static allodynia withdrawal thresholds in HFD only mice. **(H)** A trend of increasing but not statistically significant dynamic allodynia scores in HFD only mice. **(I)** Decreased static allodynia withdrawal thresholds and **(J)** increasing dynamic allodynia scores in mice receiving HFD with low-dose STZ. Static and dynamic allodynia were assessed by the von Frey and dynamic brush assays, respectively. Error bars indicate SEM. Statistics were performed by Student's t-test: *** $p < 0.001$, ** $p < 0.01$, * $p < 0.05$, ns = non-significant.

**Supplementary Fig. 2. Three-dimensional (3D) mapping of mouse foot skin.** Mouse foot skin was paraformaldehyde fixed, OCT embedded and serially sectioned. Sections were alternately stained hematoxylin and eosin (H&E), and antibody stained for PGP9.5 or S100. Six microanatomical structures were labelled in the H&E images and S100 and PGP9.5 positivity was identified in the IHC stained images using a deep learning semantic segmentation algorithm. 3D renderings of the mouse microanatomy show a high density of free nerve endings (PGP9.5$^{+}$) and Meissner corpuscles (S100$^{+}$/PGP9.5$^{+}$) in control mice, a noticeable drop in HFD (diabetic neuropathy) mice, and a total loss of Meissner corpuscles with some remaining free nerve endings in HFD + STZ (advanced diabetic neuropathy) mice.

**Supplementary Fig. 3. scRNA-seq data from WT C57BL6 mouse foot skin. (A)** UMAP projection colored by the expression of selected neurotrophins, glial cell line-derived neurotrophic factors or SIRT1.

**Supplementary Fig. 4. Leaky expression of Cre recombinase in skin of *Keratin 5 (K5)-CreER$^{T2}$* lines but no off-target effect on other tissue. (A)** Western blot analysis of skin lysates from WT, Control 2 (*K5-CreER$^{T2}$;SIRT1$^{flox/flox}$*, no tamoxifen [TAM]) and keratinocyte-specific *SIRT1* KO mice (*K5-CreER$^{T2}$;SIRT1$^{flox/flox}$*, with TAM). *SIRT1$^{flox/flox}$* mice possessed *loxP* sites flanking exon 4 of the sirtuin 1 gene. After Cre-lox recombination, a truncated SIRT1 protein product (ΔEx4) was produced. Note that in WT mice a SIRT1 isoform (ΔEx8) was also present and could not be distinguished from ΔEx4 on the Western blot. Vinculin served as a loading control. n = 2 for each group. **(B)** Leaky Cre-mediated recombination (without tamoxifen) in 31% of the foot skin basal keratinocytes compared with 76% efficiency with tamoxifen treatment in *K5-CreER$^{T2}$;R26 $^{LSL\text{-}tdTomato}$* mice. Scale bars represent 100 μm. n = 2 for each condition. **(C)** After tamoxifen-activated Cre-lox recombination, the *K5-CreER$^{T2}$;R26 $^{LSL\text{-}tdTomato}$* mice expressed tdTomato in back hairy skin in addition to foot skin (B). In comparison, the spinal cord and dorsal root ganglia (DRG) were negative for tdTomato expression. Scale bars represent 50 μm for skin and 100 μm for spinal cord and DRG.  n = 2.

**Supplementary Fig. 5. HFD-induced impaired glucose tolerance unaffected by depletion of skin keratinocyte-derived SIRT1 or oral supplementation of nicotinamide riboside (NR).** **(A)** 2-hour GTT at baseline for keratinocyte-specific *SIRT1* KO and control mice. **(B)** 2-hour GTT after 3 months of HFD for keratinocyte-specific *SIRT1* KO and control mice. **(C)** The glucose AUC. Note that all 3 groups developed similar levels of impaired glucose tolerance. ~~**(D)** 2-hour GTT and **(E)** glucose AUC of *SIRT1* KO and control mice receiving NR or vehicle (saline). **(F)** Meissner corpuscle densities and **(G)** free nerve ending densities (IENFD) of *SIRT1* KO and  control mice receiving NR or vehicle.~~ Error bars indicate SEM. Two-way ANOVA: *** $p < 0.001$, * $p < 0.05$, ns = non-significant.

**Supplementary Fig. 6. scRNA-seq data from foot skin of keratinocyte-specific *SIRT1* KO and control (*SIRT1*$^{flox/flox}$) mice. (A)** UMAP projection colored by cell type annotation (n = 3584). **(B)** UMAP plot of foot skin cells from keratinocyte-specific *SIRT1* KO and Control 1 mice (n = 2 for each condition).

**Supplementary Fig. 7. HFD-induced impaired glucose tolerance unaffected by depletion of skin keratinocyte-derived BDNF or overexpression of keratinocyte-derived SIRT1. (A)** 2-hour GTT and **(B)** glucose AUC of keratinocyte-specific *BDNF* KO and control mice at baseline and after 3 months of HFD. **(C)** 2-hour GTT and **(D)** glucose AUC of *K5-rtTA;SIRT1-TRE* mice receiving HFD only or HFD + DOX diet. Error bars indicate SEM. Statics were performed by one- or two-way ANOVA: **** $p < 0.0001$, *** $p < 0.001$, ** $p < 0.01$, ns = non-significant.

**Supplementary Fig. 8. Graphical abstract.** Diabetes causes decreased SIRT1 activity and subsequent downregulation of BDNF expression in foot skin. As a result of deficient trophic support, Meissner corpuscles and their innervating $TrkB^+$ Aβ axons degenerate. Retrograde degeneration of peripheral axons may trigger hyperexcitation of $TrkB^+$ mechanosensory neurons, resulting in mechanical allodynia.

### SUPPLEMENTARY FIG. 1

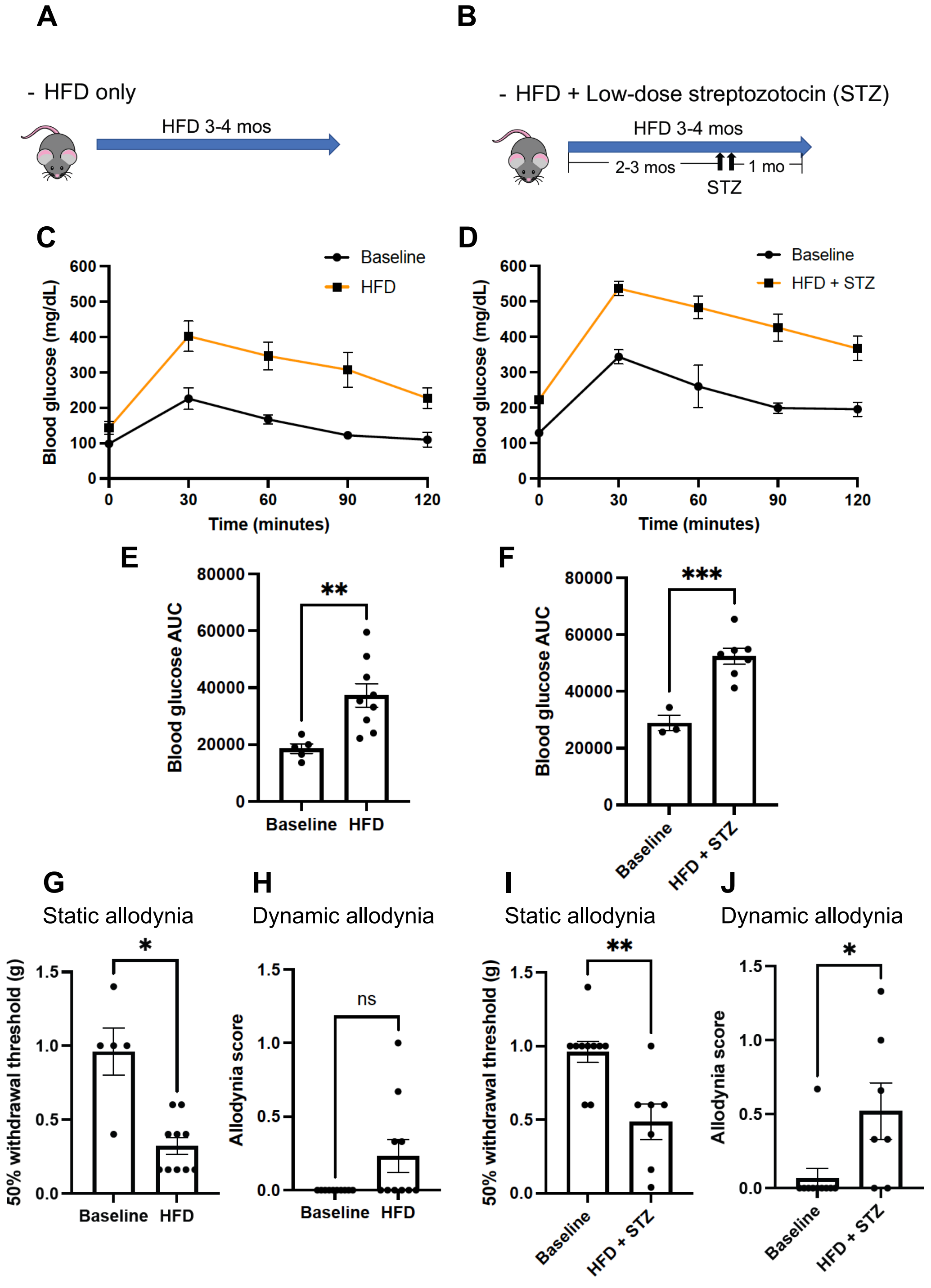

### SUPPLEMENTARY FIGURE 2

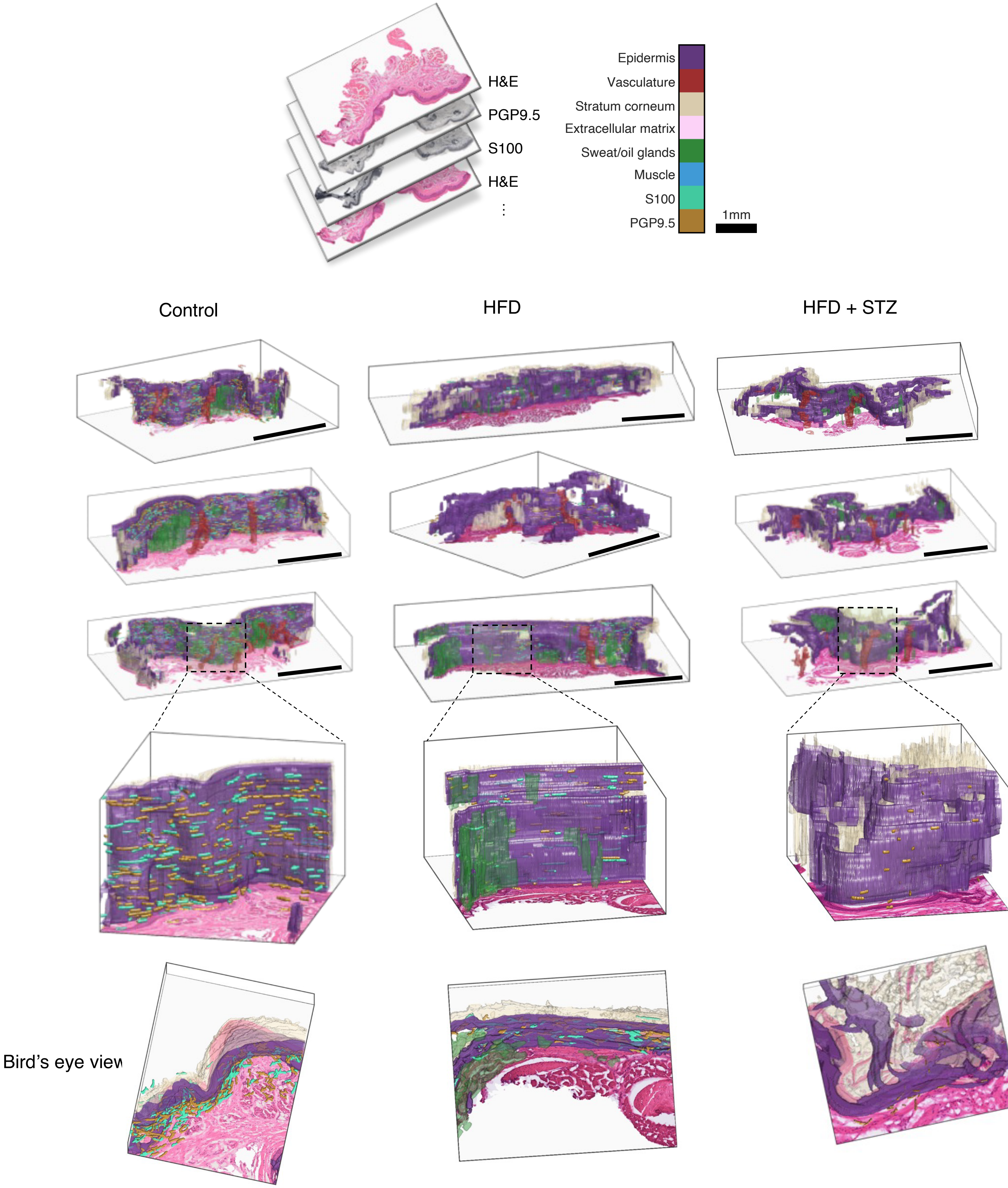

### SUPPLEMENTARY FIG. 3

**A**

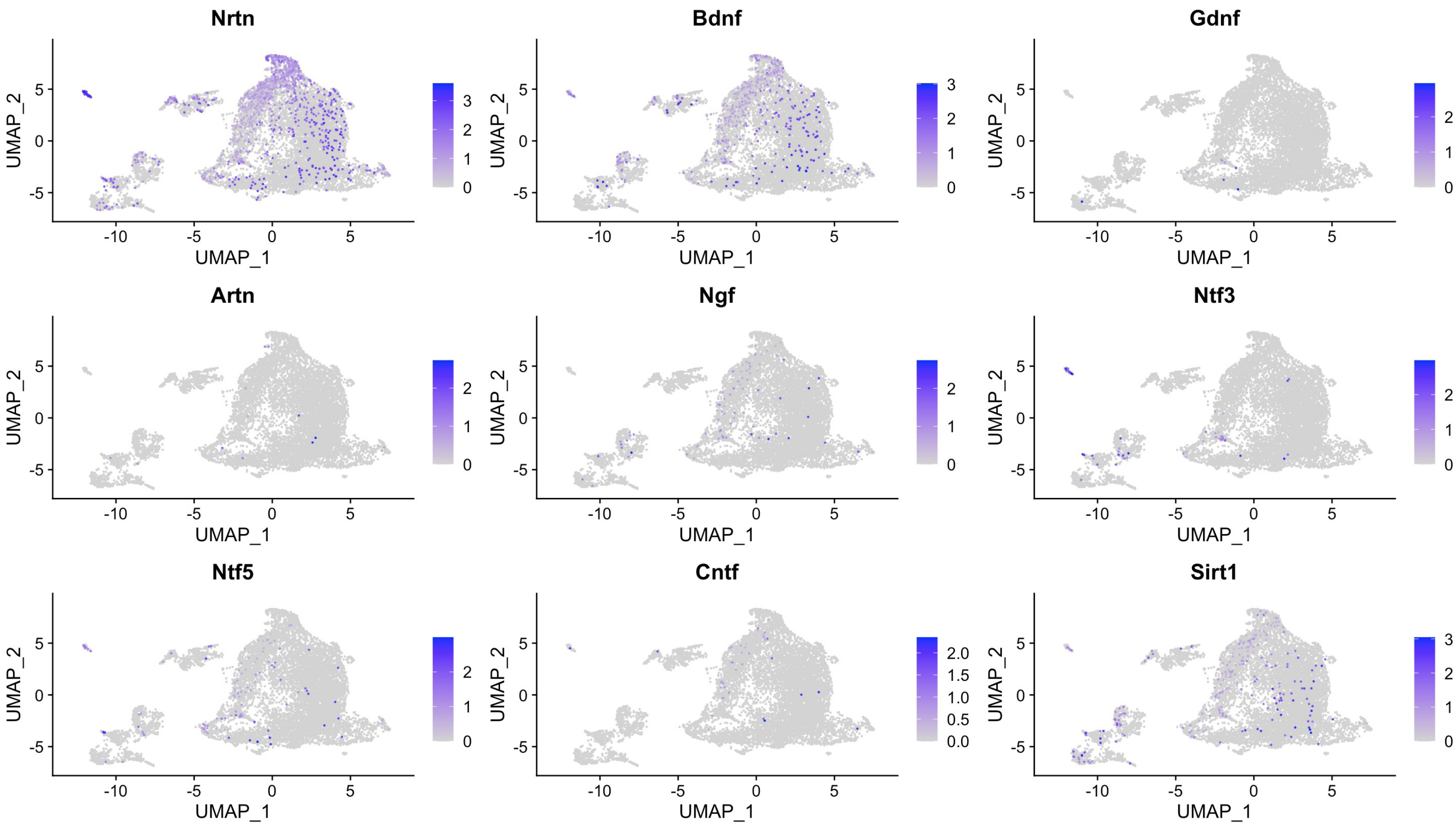

### SUPPLEMENTARY FIG. 4

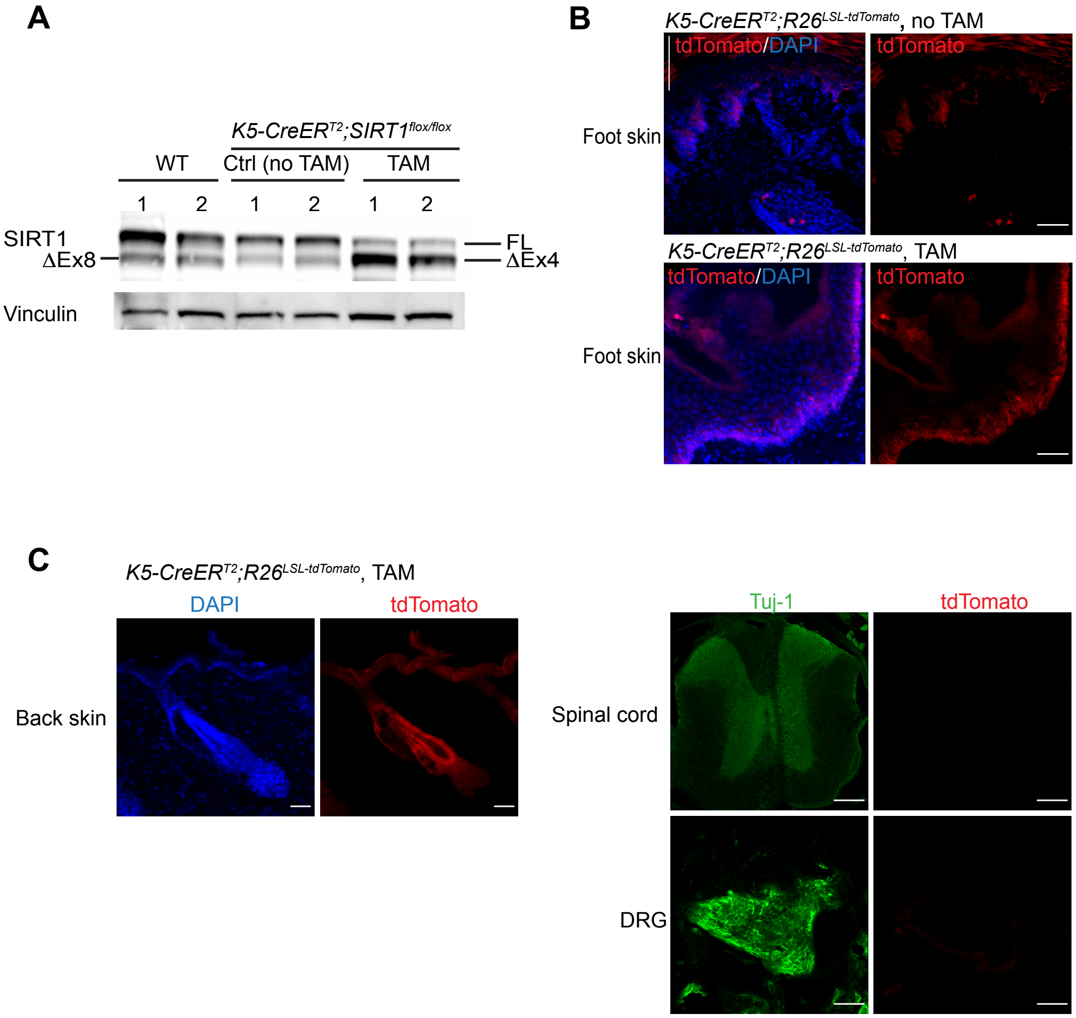

### SUPPLEMENTARY FIG. 5

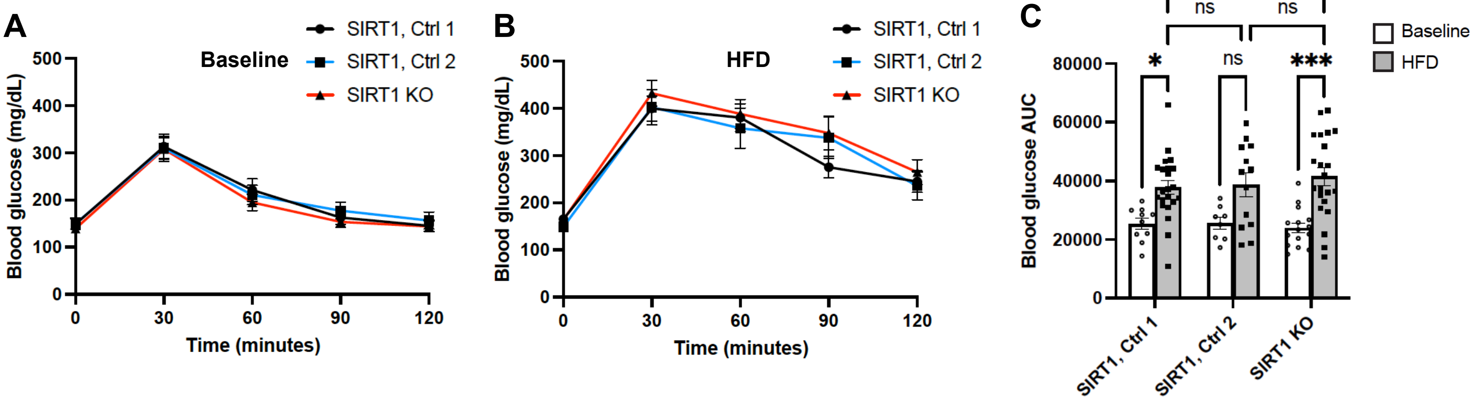

### SUPPLEMENTARY FIG. 6

A

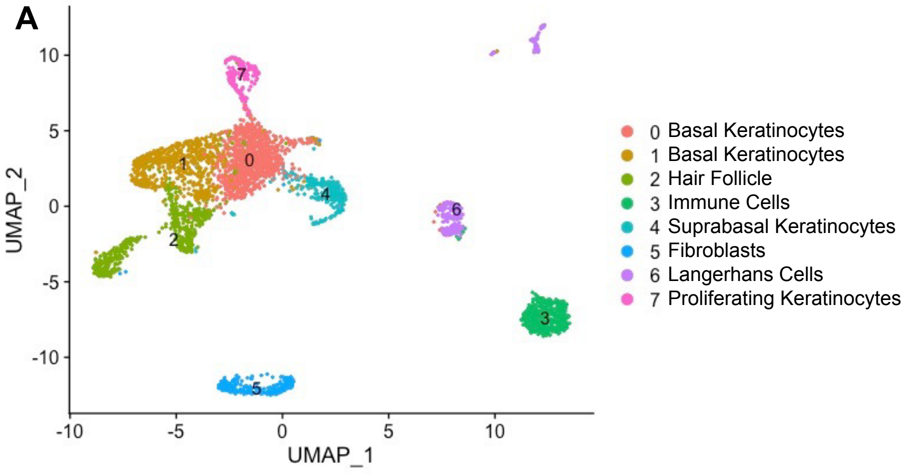


B

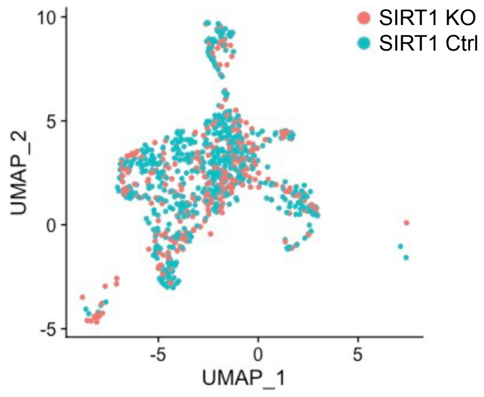

### SUPPLEMENTARY FIG. 7

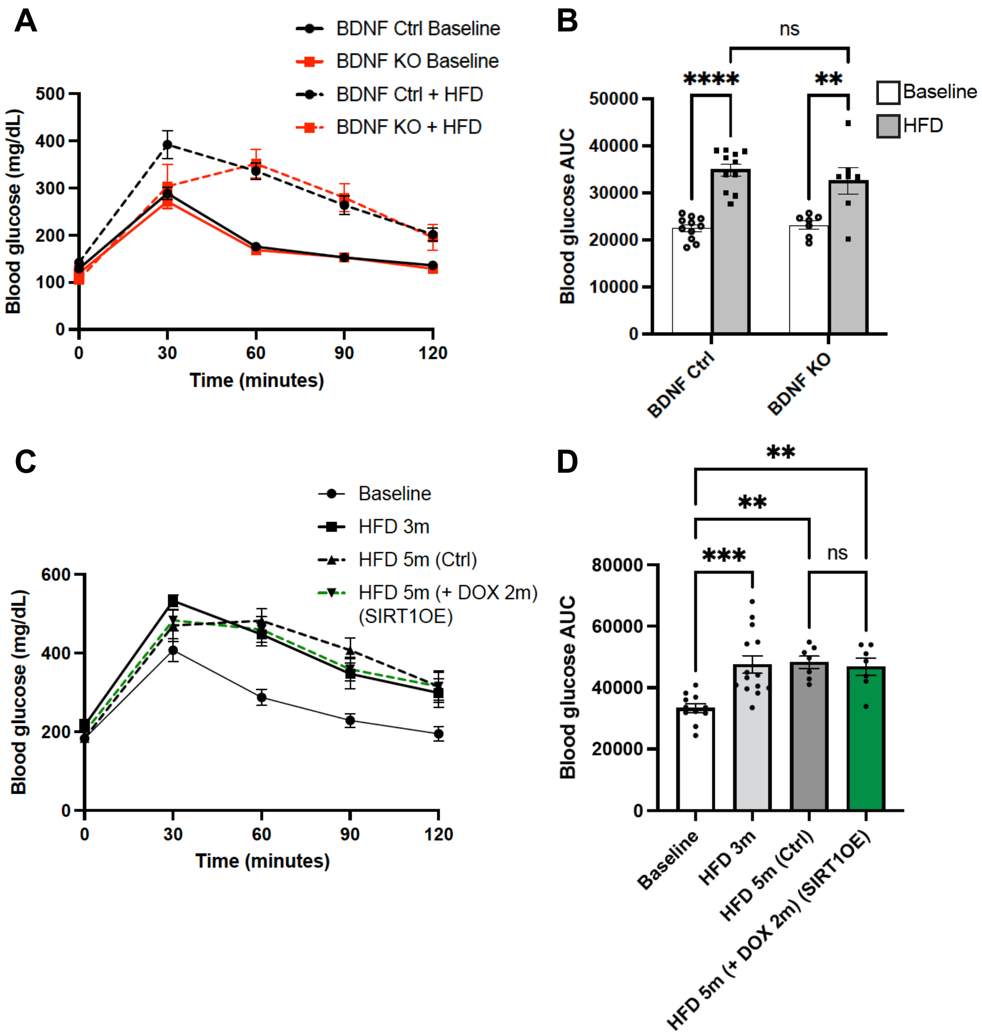

### SUPPLEMENTARY FIGURE 8

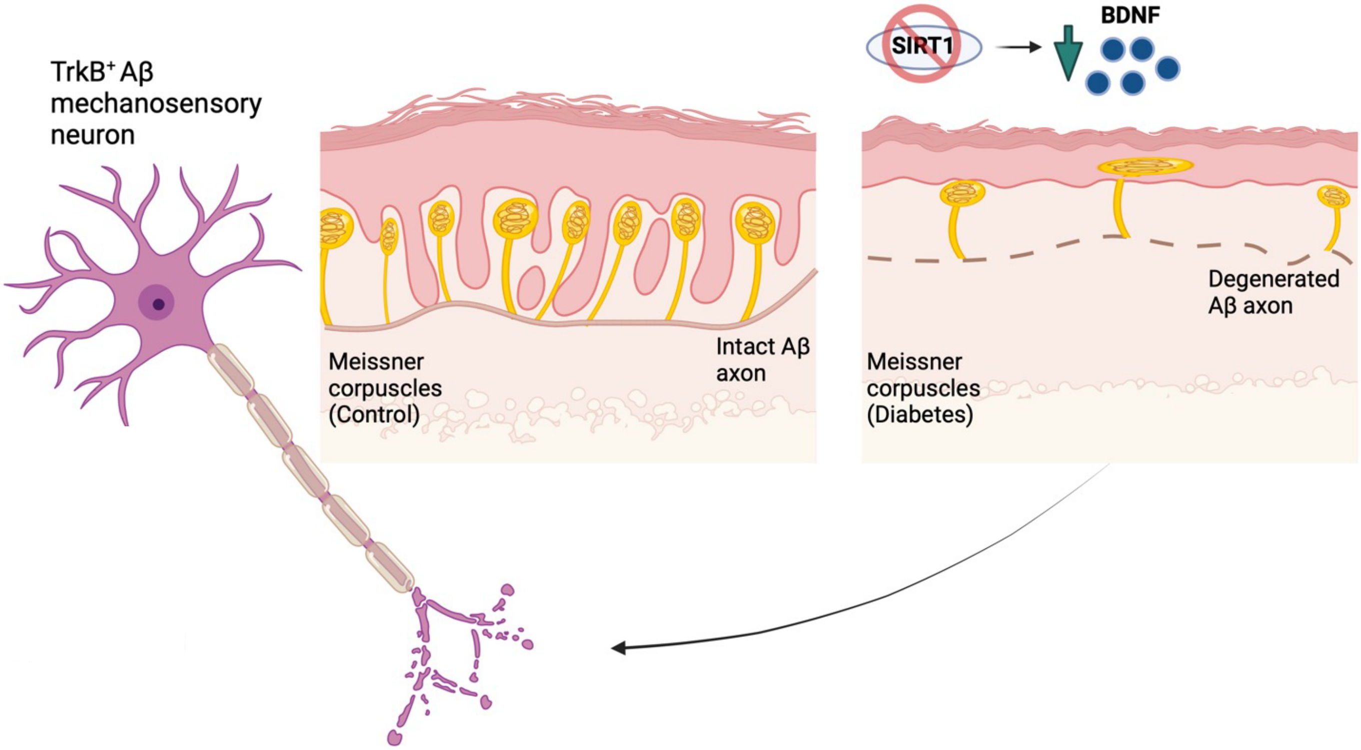
